# Alzheimer’s Disease Risk Allele *APOE4* Interacts with Arsenic Exposure to Drive Microglial Dysfunction

**DOI:** 10.64898/2026.05.09.723490

**Authors:** Alex J. Marchi, Ashley M. Brooks, Elizabeta Gjoneska

**Affiliations:** Neuroepigenomics Group, Neurobiology Laboratory, National Institute of Environmental Health Sciences; Genetics and Genomics Department, North Carolina State University; Integrative Bioinformatics Support Group, National Institute of Environmental Health Sciences

## Abstract

Alzheimer’s disease (AD) is influenced by both genetic risk and environmental exposures, but how these factors interact in human microglia remains unclear. Here, we investigate whether the late-onset AD risk allele *APOE4* impacts microglial vulnerability to arsenite exposure.

To that end, we used CRISPR/Cas9 to generate an isogenic *APOE4*^+/+^ iPSC-derived transcription factor-induced microglia-like cells (iTFM). We demonstrate that *APOE4*^+/+^ iTFM exhibit decreased survival following arsenite exposure, as evidenced by a lower LC^50^ compared to *APOE3*^+/+^ controls. Transcriptomic profiling identified arsenite concentration as the primary driver of gene expression changes, while genotype contributed a secondary, distinct component of the response. Weighted gene co-expression network analysis revealed genotype-dependent modules enriched for phagocytic and oxidative stress pathways, including KEAP1-NFE2L2 signaling. These transcriptomic changes were further supported by functional assays. *APOE4*^+/+^ iTFM had a high proportion of phagocytic cells and altered mitochondrial phenotypes including increased mitochondrial mass, reduced membrane potential, and reduced superoxide production, all of which were further perturbed by low dose arsenite exposure.

These results support a gene-environment interaction-dependent increase in microglial vulnerability via reshaping of transcriptional and functional stress responses, and provide a human cell-based framework for studying environmentally mediated microglial contributions to AD.

## INTRODUCTION

Dementia is the 7^th^ leading cause of death worldwide with approximately 10 million new cases diagnosed each year. The most common form of dementia is Alzheimer’s Disease (AD), which accounts for 60-70% of all cases^1^. AD is a progressive neurodegenerative disorder characterized by a gradual decline in memory and cognition accompanied by pathological protein aggregation, neuronal and synaptic loss, and chronic neuroinflammation^2–5^. While AD is classically defined by the formation of extracellular amyloid-β (Aβ) plaques and intracellular neurofibrillary tangles, major aspects of its pathogenesis remain unresolved. In particular, important gaps persist in understanding of the triggers that initiate Aβ and tau pathology as well as how these factors interact with neuroinflammatory, genetic, and cell-type-specific mechanisms to drive selective vulnerability and disease progression.

Among these unresolved mechanisms, neuroinflammation has emerged as a critical contributor to AD pathology. Microglia, the resident macrophages of the central nervous system (CNS), are key regulators of immune surveillance, phagocytosis, synaptic remodeling, and responses to pathological stress, placing them at the center of disease-relevant processes in AD^6^. In a healthy brain, microglia preserve tissue homeostasis. However, in the context of AD, sustained microglial activation and dysregulation are thought to influence Aβ and tau pathology by altering clearance capacity, inflammatory signaling, and neuronal homeostasis^7–10^. These findings have heightened interest in the factors that shape microglial state and vulnerability, particularly the AD-associated risk allele *APOE4*. Apolipoprotein E (*APOE*) encodes apoE, a key lipid transport protein involved in cholesterol homeostasis, membrane remodeling, and receptor-mediated cellular responses in both peripheral tissues and the CNS^11^. *APOE4*^+/+^ is the strongest common genetic risk factor for late onset AD. In microglia, *APOE4*^+/+^ has been associated with activation of inflammatory genes and altered Aβ uptake, phagocytosis, and motility^12,13^. However, genetic risk alone may not fully account for disease-associated microglial dysfunction. Microglia are highly responsive to their environment, making it essential to understand how external exposures interact with genetic susceptibility to shape their functional state and stress response. Among such exposures, arsenic is of particular interest due to its known ability to disrupt oxidative balance, mitochondrial function, and inflammatory signaling.

Arsenic is a well-documented, naturally occurring metalloid toxicant that is distributed throughout soil, water, rocks, and air. Unlike other heavy metal toxicants, ingested arsenic has a 90% absorption rate in the gastrointestinal tract^14,15^. The primary form of human arsenic exposure is through contaminated drinking water^16–18^ and therefore the US Environmental Protection Agency regulates arsenic in drinking water with a maximum level of 10 parts per billion (ppb). Nevertheless, arsenic levels exceeding this threshold have been detected in at least 7% of wells sampled across the US^19^. For example, in North Carolina, arsenic levels as high as 806 ppb have been detected in unregulated, private wells^20^. Arsenic has been shown to pass through the blood brain barrier in mice^21–23^ and levels correlate with increased AD rates^24,25^. Furthermore, higher levels of arsenic have been found in post mortem brains of AD patients^26^, warranting further investigation into arsenic as a potential contributor to late onset AD. In microglia, arsenic induces microgliosis, inflammatory signaling via TNFα, IL-1β, and CD68, and neurodegeneration in mice^23,27^, however there is a scarcity of research performed in human models. This is especially important due to the differences in the arsenic metabolism between rodents and humans. For example, mice eliminate arsenic much more rapidly through urinary excretion of dimethylarsinic acid compared to humans, who do so via monomethylarsonic acid^16,28,29^. What remains unclear is the extent to which *APOE4* mediated genetic susceptibility interacts with arsenic exposure to drive microglial dysfunction, thereby contributing to late-onset AD pathology.

To better understand the relationship between environmental exposures and AD-associated genetic risk in microglia, we utilized transcription factor inducible microglia-like cells (iTFM), which enable us to isolate the cell-type specific effect of the interaction. These cells are rapidly differentiated from KOLF2.1J induced pluripotent stem cells (iPSCs) within 8 days and faithfully recapitulate lineage-specific markers of primary microglia^30–32^. Using this system, we generated an isogenic *APOE4* model and exposed it to various concentrations of arsenic to investigate interactions between microglial AD genetic susceptibility and their response to environmental stressors.

## RESULTS

### KOLF2.1J-iTFM Recapitulate Molecular Signatures of Microglia

We first set out to validate whether the KOLF2.1J iTFM faithfully recapitulate the molecular signatures of microglia as other previously described iTFM^31^. To that end, we differentiated KOLF2.1J iPSCs and observed that differentiated iTFM morphologically resembled microglia starting at day 8 (Figure 1A). Of note, small groups of differentiated non-microglial cells constituting < 5% of the population were occasionally detected.

**Figure 1:**
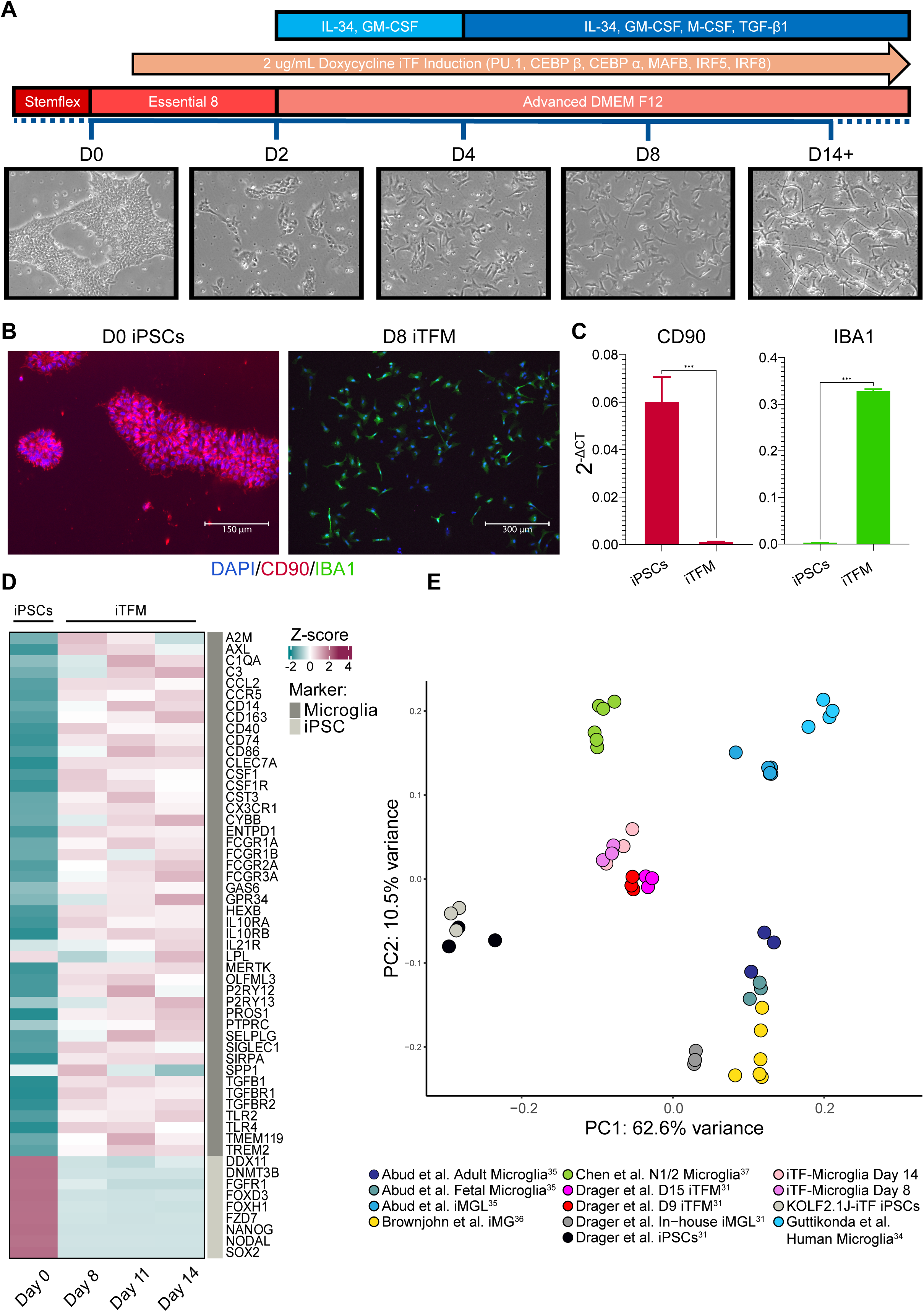
(A) iTFM differentiation protocol depicting cytokines added (blue), transcription factors induced (orange), and media used (red/pink) with reference brightfield images depicting cell morphology. (B) Immunocytochemistry showing DAPI (blue), iPSC marker CD90 (red), and the microglial marker IBA1 (green) in undifferentiated iPSCs and D8 iTFM. (C) Gene expression levels of CD90 and IBA1 measured by RT-qPCR. (D) Heatmap showing averaged expression (n=3) of iPSC and microglia marker genes in iPSCs and iTFM on day 8, 11, and 14 of differentiation. (E) PCA of gene expression data from our iTFM (pink and purple) and other relevant iTFM, iMGL, and microglia samples. Colors represent different cell types.

Immunocytochemistry and RT-qPCR revealed a loss of the iPSC marker CD90 and gain of microglial marker IBA1 by day 8 (students t-test, p = 0.0006, <0.0001 respectively) (Figure 1B, 1C). Downregulation of iPSC markers alongside an increase of microglial markers was also confirmed by RNA sequencing (RNA-seq) of day 0 iPSCs, day 8, and day 14 iTFM (Figure 1D). Moreover, the iTFM transcriptomic profile was nearly identical to that of other iTF-derived microglia and similar to iPSC-derived microglia produced using different differentiation methods (Figure 1E).

### Generation of an Isogenic *APOE4*^+/+^ iPSC line

To investigate the effects of arsenic on AD microglia, we generated an isogenic *APOE4*^+/+^ mutation in the parental *APOE3*^+/+^ (wild-type) KOLF2.1J-iTF iPSC cell line using CRISPR/Cas9 mediated homology directed repair. The *APOE4*^+/+^ Cys112Arg missense mutation was verified via sanger sequencing of expanded monoclones (Supplementary Figure 1A). Whole genome sequencing of both the parental cell line and two *APOE4*^+/+^ monoclones revealed a comparable number of mutations with greater >92% sites shared and a similar distribution of predicted functional consequences (Supplementary Figure 1B, 1C). We also observed similar gene expression profiles between *APOE4*^+/+^ monoclones by RNA-seq (Supplementary Figure 1D).

### *APOE4*^+/+^ iTFM Display Increased Sensitivity to Arsenite Exposure

To evaluate the impact of *APOE4*^+/+^ on iTFM vulnerability to arsenic, we exposed day 8 iTFM to varying concentrations of arsenite, the main inorganic form of arsenic^33^, for 72 hours. Significant genotype dependent differences in normalized survival were observed at 0.1, 1, 5, 10 and 50 µM (students t-test, p = 0.003, 0.0029, 0.0045, 0.0139, 0.0101 respectively) (Figure 2A). Consistent with this, *APOE4*^+/+^ iTFM displayed a lower arsenite LC^50^ than *APOE3*^+/+^ cells (6.63 vs 8.13 µM, respectively).

**Figure 2:**
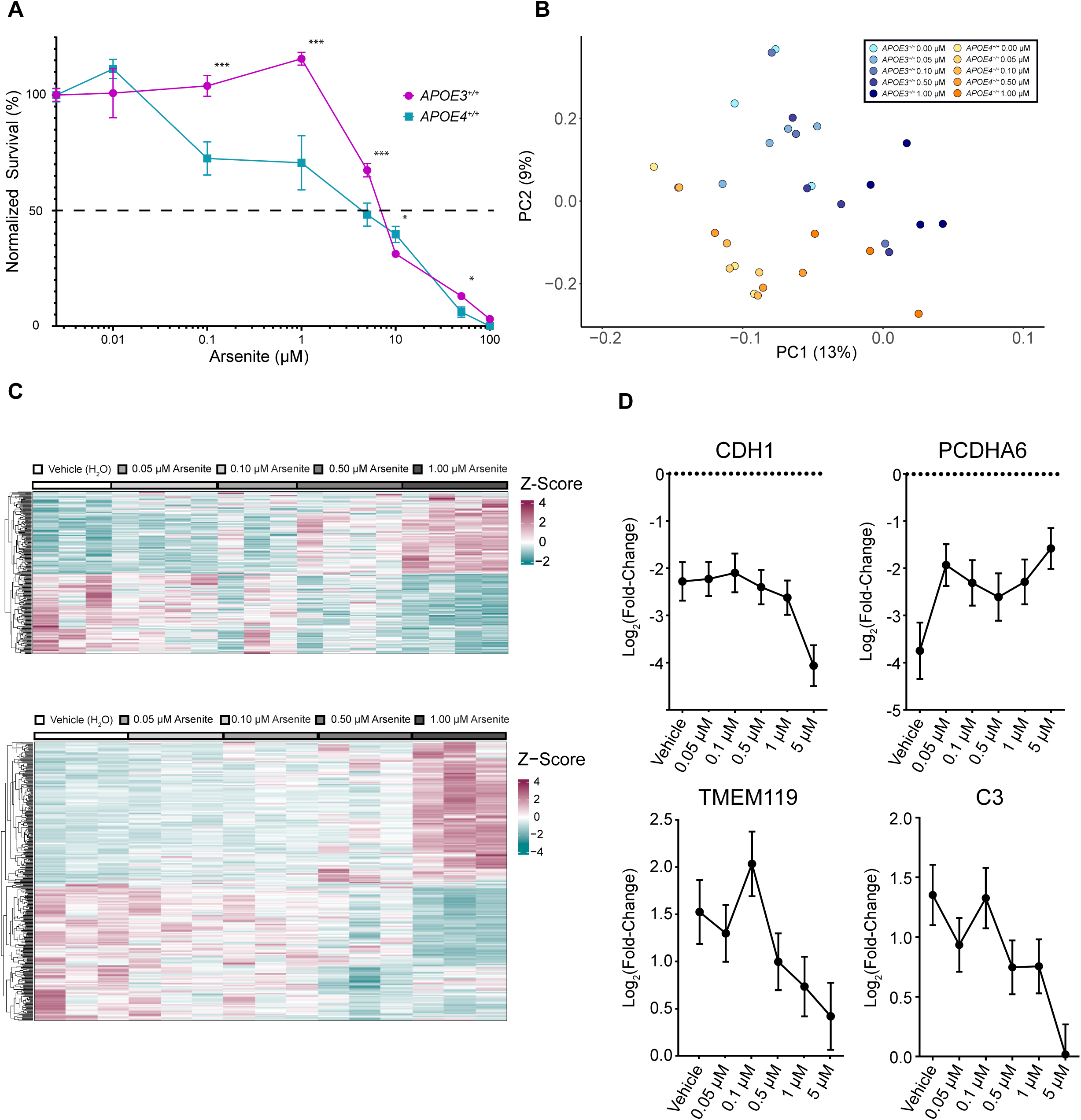
(A) Normalized survival of 72-hour arsenite treated APOE3+/+ and APOE4+/+ iTFM. *, **, *** = p <0.05, 0.005, and 0.001 respectively. APOE3+/+ LC50 = 8.13 μM, APOE4+/+ LC50 = 6.63 μM. (B) PCA of 72-hour arsenite treated APOE3+/+ (blue) and APOE4+/+ (yellow) iTFM. Each dot represents an independent biological sample responding to increasing concentration of arsenite (light to dark). (C) Heatmaps of APOE3+/+ (top) and APOE4+/+ (bottom) differentially expressed genes in response to increasing arsenite concentration. (D) Plots showing Log2(Fold-Change) of specific microglia genes in APOE4+/+ iTFM relative to APOE3+/+ at each arsenite concentration.

To determine how the *APOE4*^+/+^ allele influences gene expression in the context of arsenite exposure, we examined transcriptional profiles of day 8 differentiated *APOE3*^+/+^ and *APOE4*^+/+^ iTFM in response to increasing concentrations of arsenite. Principle component analysis (PCA) revealed that variation along PC1 was primarily driven by arsenite concentration, while variation along PC2 reflected a separation by genotype (Figure 2B). To investigate this further, an RNA-seq analysis was performed to identify significant differentially expressed genes (Supplementary Figure 4). We observe minor gene expression changes at low arsenite concentrations (0.05 and 0.1 μM) in both *APOE3*^+/+^ and *APOE4*^+/+^ iTFM that becomes more significant as arsenite exposure increases (Figure 2C, Supplementary Figure 2, 3). Moreover, pronounced gene expression differences were observed between *APOE3*^+/+^ and *APOE4*^+/+^ iTFM at each arsenite concentration examined (Supplementary Figure 4). Ingenuity Pathway Analysis (IPA) identified altered molecular pathways, biological processes, and upstream regulators (Supplementary Figure 5). Among the most significantly differentially expressed genes across arsenite concentrations were CDH1, PCDHA6, TMEM119, and C3 (Figure 2D).

### *APOE4*^+/+^ Interacts with Arsenite Exposure to Drive Microglia Dysfunction

To identify coordinated transcriptional responses associated with arsenite exposure, we ran weighted gene co-expression network analysis (WGCNA) across all arsenite treated RNAseq datasets and examined how module behavior changed with increasing arsenite concentrations in *APOE3*^+/+^ and *APOE4*^+/+^ iTFM. This approach allowed us to define concentration-associated gene networks and compare their robustness, directionality, and magnitude between genotypes. PCA revealed clear separation by arsenite concentration along PC1 and genotype along PC2 (Supplementary Figure 6). We further identified multiple modules with genotype-specific transcriptional responses to increasing arsenite exposure (Figure 3A, 4A, Supplementary Figure 8, Supplementary Table 1). Subsequent ingenuity pathway analysis of these gene sets identified significant differences in pathways involved in phagocytosis (“magenta” module) and oxidative stress response (“black” module), such as the KEAP1-NFE2L2 pathway, between *APOE3*^+/+^ and *APOE4*^+/+^ iTFM (Figure 3B, 4B).

**Figure 3:**
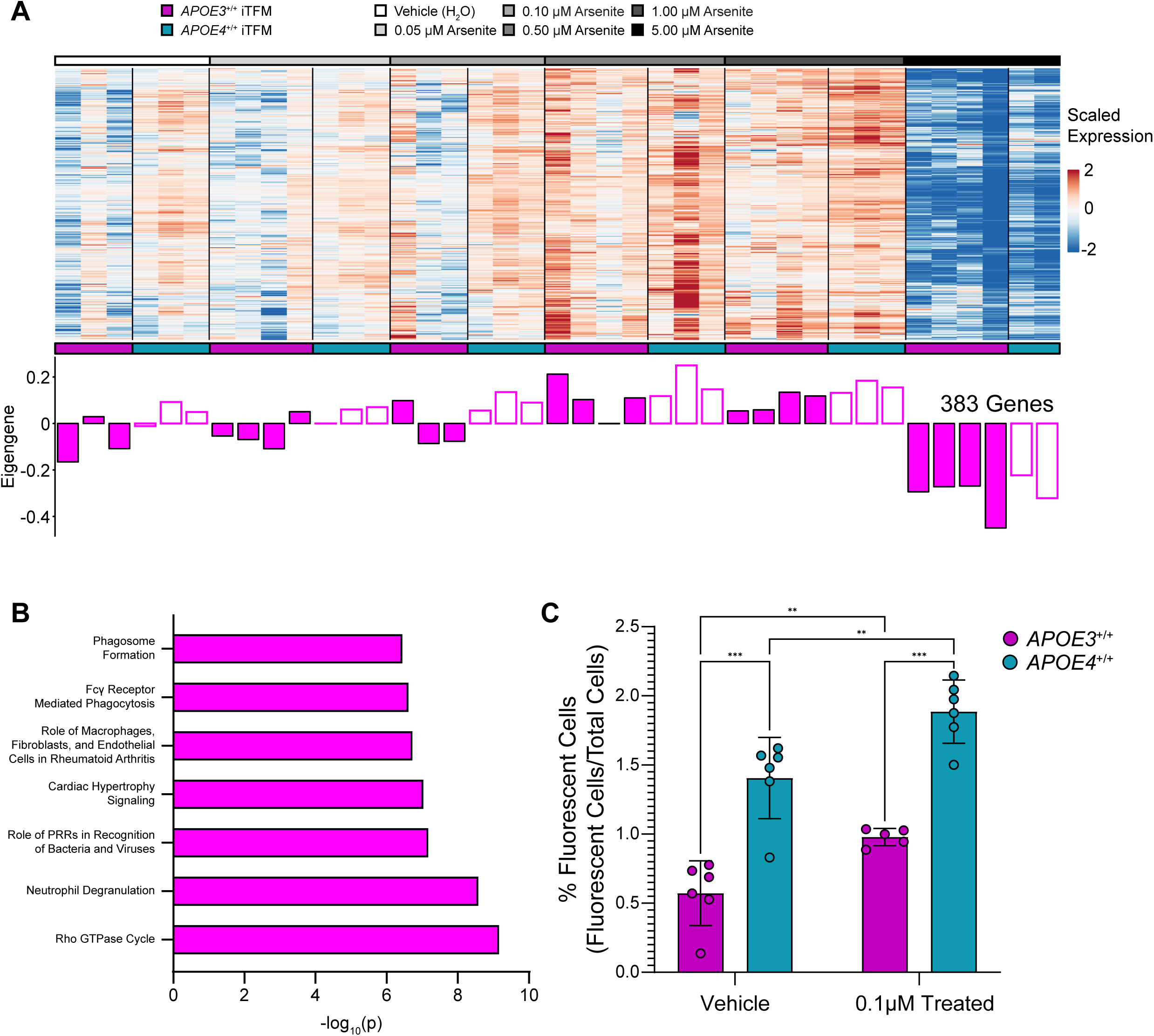
(A) Co-expression analysis showing relative gene expression changes from the “magenta” gene module between APOE3+/+ and APOE4+/+ iTFM after 72 hrs exposure to increasing arsenite concentrations. (B) Pathways enriched in the “magenta” gene module organized by −log10(p). (C) Percent iTFM that phagocytosed one or more fluorescent zymo beads measured by flow cytometry. Statistical significance assessed by two-way ANOVA followed by Bonferroni post hoc testing. *, **, and *** correspond to p values <0.05, 0.005, 0.0001 respectively.

**Figure 4:**
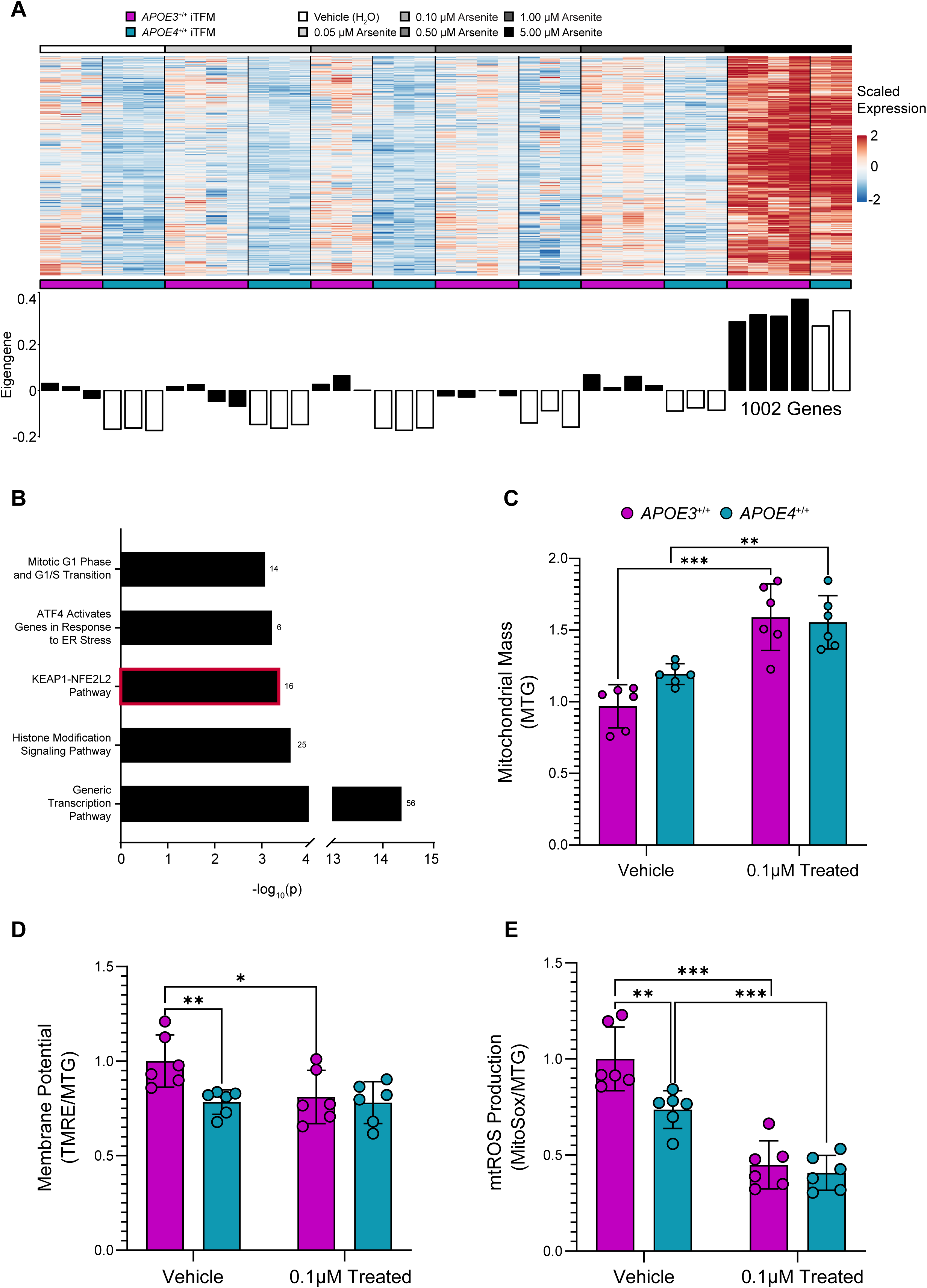
(A) Co-expression analysis showing the relative gene expression changes from the “black” gene module between APOE3+/+ and APOE4+/+ iTFM after 72 hrs exposure to increasing arsenite concentrations. Pathways enriched in the “black” gene module organized by −log10(p). (C) Mitochondrial mass of iTFM measured by mitotracker green (MTG) staining. (D) Mitochondrial membrane potential measured by Tetramethylrhodamine ethyl ester (TMRE) normalized by MTG. (E) Mitochondria superoxide production measured by mitosox red normalized to MTG. in (C-E) APOE3+/+ (purple) and APOE4+/+ (teal) iTFM. Statistical significance assessed by two-way ANOVA followed by Bonferroni post hoc testing. *, **, and *** correspond to p values <0.05, 0.005, 0.0001 respectively.

We further validated transcriptomic differences observed in the “magenta” module (Figure 3A, 3B) by quantifying the ability of arsenite treated *APOE3*^+/+^ and *APOE4*^+/+^ iTFM to phagocytose fluorescent 1 μm latex beads. These beads can be internalized, but not degraded, thereby enabling quantification of both the proportion of phagocytically active iTFM and their phagocytic capacity. When assessing phagocytic capacity through mean fluorescent intensity, we observed no differences between genotype or exposure (Supplementary Figure 7) (two-way ANOVA, Bonferroni corrected). However, consistent with the pathway enrichment identified by WGCNA and IPA, we found a significantly greater proportion of phagocytic *APOE4^+/+^* iTFM compared to *APOE3^+/+^* cells under both vehicle and 0.1 µM arsenite (two-way ANOVA, Bonferroni corrected; p = <0.0001 for both comparisons). Notably, exposure to 0.1 µM arsenite (13 ppb) significantly increased the proportion of phagocytic iTFM relative to vehicle treated controls, especially in *APOE4^+/+^* populations (Figure 3C) (two-way ANOVA, Bonferroni corrected; p = .0082, p = .0017 respectively), suggesting that *APOE4^+/+^* iTFM are more likely to engage in phagocytosis and this is further exacerbated by arsenite treatment.

To validate transcriptomic changes observed in the “black” module (Figure 4A, 4B), we performed functional assays quantifying mitochondrial dysfunction by measuring mitochondrial mass, membrane potential and superoxide production. We observed increases in mitochondrial mass in both genotypes upon exposure to 0.1 µM (13ppb) arsenite. Untreated *APOE4*^+/+^ had a higher baseline level (Figure 4C). Quantification of mitochondrial membrane potential indicated that exposure to as little as 0.1 µM (13ppb) arsenite is sufficient to reduce membrane potential of *APOE3*^+/+^ iTFM to levels of *APOE4*^+/+^ iTFM (Figure 4D). Mitochondria superoxide production (mtROS) of untreated *APOE4*^+/+^ was also lower than *APOE3*^+/+^ controls, and exposure to 0.1 µM (13ppb) arsenite further reduced mtROS production irrespective of genotype (Figure 4E). Together, these results suggest that even 13 ppb arsenic, just above the EPA limit, is sufficient to cause significant mitochondrial changes in iTFM.

## DISCUSSION

In this study, we validate a KOLF2.1J derived iTFM model and generate an isogenic *APOE4*^+/+^ iTFM cell line using CRISPR/Cas9, which minimizes background genetic variation allowing observed differences to be more confidently linked to genotype. Here we report that *APOE4*^+/+^ modifies the iTFM response to arsenite exposure as indicated by reduced viability, and altered transcriptional state, mitochondrial function, and phagocytic behavior.

The KOLF2.1J iTFM system consistently and rapidly produced microglial like cells characterized by the loss of pluripotency markers, gain of microglial markers, and transcriptomic profiles similar to those of previously described iTFM^31,34–37^. While small clusters of non-iTFM cells were occasionally present, they did not appear to meaningfully distort the overall signal. Together, these findings support the KOLF2.1J iTFM system as a reliable human microglial model for toxicology and AD-risk studies.

When exposed to increasing concentrations of arsenite, *APOE4*^+/+^ iTFMs exhibited reduced survival as measured by the lower LC^50^ (6.63 μM) compared to *APOE3*^+/+^ iTFMs (8.13 μM), with significant viability loss beginning at 0.1 μM arsenite exposure. These results indicate that *APOE4*^+/+^ iTFMs are more vulnerable to arsenite toxicity and suggest that *APOE4*^+/+^ associated microglia may be more susceptible to environmentally relevant toxicant exposure. Because microglia are critical mediators of immune surveillance and homeostasis in the central nervous system^6^, arsenite induced reduction in microglial viability could compromise the CNS immune defense and increase vulnerability to other neurological insults. Whether this heightened vulnerability is specific to arsenite or reflects a broader susceptibility to environmental toxicants remains to be determined.

Transcriptional profiling across increasing arsenite concentrations revealed that the primary transcriptional variation was driven by exposure level, whereas secondary variation was associated with genotype. These findings support a gene by environment interaction in which arsenite acts as the dominant perturbation and *APOE4*^+/+^ modifies how iTFMs respond and adapt to that insult. Although DEG analysis revealed substantial transcriptional differences across genotype and arsenite concentration, no significant DEGs were detected in either *APOE3*^+/+^ or *APOE4*^+/+^ iTFM at 0.05 or 0.1 μM arsenite relative to vehicle controls. This suggests that low dose arsenite exposure may not produce large transcriptomic changes detectable by conventional DEG thresholds. Instead, the response may involve subtle, but coordinated, shifts at the pathway level^38,39^, or post-transcriptional and metabolic adaptation that alter cellular function without markedly changing mRNA abundance^40^. This interpretation is consistent with evidence that environmentally relevant low dose arsenite can elicit subtle, nonlinear biological responses^41^. WGCNA demonstrated coordinated transcriptional responses associated with arsenite exposure across all RNA-seq datasets. Specifically, we identified co-expression modules whose relationship with arsenite concentration differed by genotype, indicating that the *APOE4*^+/+^ allele shapes the organization of the transcriptional response to exposure. IPA of these modules revealed differential regulation of phagocytic pathways and the KEAP1-NFE2L2 oxidative stress pathway. Together, these findings support a model in which *APOE4*^+/+^ not only affects the expression of individual genes but also impacts broader stress response networks that shape microglial adaptation and response to environmental toxicants.

The enrichment of phagocytosis-related pathways is consistent with prior studies implicating *APOE4*^+/+^ in altered microglial phagocytic function; however, the literature is mixed, which may suggest that *APOE4*^+/+^ modifies the regulation and context of phagocytosis rather than uniformly increasing or decreasing it^12,13,42–45^. The increased phagocytic engagement may be beneficial, however excessive or dysregulated phagocytosis can reflect a reactive microglial state relevant to neurodegeneration^46,47^. Together, these results suggest that *APOE4*^+/+^ alters microglial phagocytic function^48,49^ and supports a model in which *APOE4*^+/+^ shifts microglia towards a more phagocytically prone state under arsenite exposure. The KEAP1-NFE2L2 finding is biologically concordant with the well-established oxidative stress response to arsenite^50,51^. Increased mitochondrial mass of untreated *APOE4*^+/+^ iTFM relative to *APOE3*^+/+^ controls which was further increased upon arsenite exposure in both genotypes, may reflect compensation for mitochondrial stress or impaired clearance of damaged mitochondria. This is consistent with prior reports of disrupted mitophagy in *APOE4*^+/+^ microglia^44^. *APOE4*^+/+^ iTFM also showed reduced mitochondrial membrane potential at baseline, resembling the phenotype induced by arsenite exposure. Reduced membrane potential is consistent with mitochondrial depolarization, disruption of the proton gradient, impaired oxidative phosphorylation, and increased cellular stress^52^. Finally, the reduced baseline mtROS observed in untreated *APOE4*^+/+^ iTFMs, which was further reduced in both genotypes following arsenite exposure, can indicate enhanced antioxidant defense. However, this reduction occurs alongside a drop in mitochondrial membrane potential and an increase in mitochondrial mass. In this context, the pattern points to mitochondrial dysfunction or the buildup of less functional mitochondria, rather than improved mitochondrial efficiency^44,53–55^. Together, these findings suggest that *APOE4*^+/+^ iTFM exist in a functionally distinct state that may predispose them to maladaptive responses under toxic stress where even low dose arsenite is sufficient to further perturb microglia homeostasis.

Future studies should determine whether the increased vulnerability observed in *APOE4*^+/+^ iTFM is specific to arsenite or extends to other environmentally relevant toxicants, including additional metals, air pollutants, and pesticides. Extending this work into more complex systems such as co-cultures, brain organoids, or xenograft models will help define how *APOE4*^+/+^ microglia influence surrounding tissue in the context of toxicant exposure. Chronic low dose exposures, broader concentration ranges, and higher resolution approaches such as single-cell transcriptomics may further clarify whether *APOE4*^+/+^ promotes distinct microglial states under environmental stress. Additionally, investigating the *APOE2*^+/+^ variant of *APOE* in the context of arsenite exposure will also be informative. *APOE2*^+/+^ has been shown to act as a protective factor against AD, reducing risk by up to 40% and delaying disease onset, therefore suggesting it could mitigate arsenite-associated effects^56^.

Overall, our results highlight a critical role for genetic variation in modulating the microglial response to environmental toxicants and support a model in which *APOE4*^+/+^ confers susceptibility to arsenite exposure by reshaping transcriptional and functional stress responses in microglia. Finally, our study establishes iTFM as a powerful platform for investigating gene-environment interactions, enabling precise isolation of cell-type specific effects within isogenic backgrounds thereby minimizing cellular heterogeneity and genetic variability.

## MATERIALS AND METHODS

### iPSC Cell Culture

KOLF2.1J-iTF iPSCs were cultured in StemFlex Basal medium (Gibco, Cat. No. A3349401) on tissue culture plates coated with Synthemax II-SC (Corning, Cat. No. 3535) with media being replaced every two days. Once 80-90% confluent, iPSCs were washed with DPBS (Gibco, Cat. No. 14190144) and lifted by incubating with StemPro Accutase cell dissociation reagent (Gibco, Cat. No. A1110501) for 7 minutes at 37°C. Accutase was neutralized by diluting 2:3 in StemFlex, cells were pelleted at 300g for 5 minutes, supernatant was aspirated, and the cell pellet was resuspended in StemFlex containing 1X RevitaCell (Gibco, Cat. No. A2644501) ROCK inhibitor. Cells were counted using Trypan blue (Gibco, Cat. No. 15250061) method in a Countless^TM^ 3 FL Automated Cell Counter (Invitrogen, Cat. No. AMQAF2000) with Countess™ Cell Counting Chamber Slides and Holder (Invitrogen, Cat. No. C10228). The desired number of cells were then passaged onto Synthemax coated plates.

### Generation of APOE4+/+ Model via CRISPR/Cas9 Editing

CRISPR/Cas9 editing was performed in accordance with previously described methods^57^ using Alt-R S.p. HiFi Cas9 Nuclease V3 (IDT, Cat. No. 1081058). Previously described modified sgRNA (Synthego; Sequence: 5’-CCUCGCCGCGGUACUGCACC-3’) and previously described^12^ Alt-R HDR modified ssODN repair template (IDT; Sequence: 5’- GAGGAGACGCGGGCACGGCTGTCCAAGGAGCTGCAGGCGGCGCAGGCCCGGCTGGCGCGGACATGGAGGACGTGCGCGGCCGGCTGGTGCAGTACCGCGGCGAGGT GCAGGCCATGCTCGGCCAGAGCACCGAGGAGCTGCGGGTGCGCCTCGCCTCCCACCTGCGCAAGCTGCGTAAG-3’) were used to generate an *APOE4*^+/+^ mutant line via CRISPR/Cas9 mediated homology directed repair. KOLF2.1J-iTF cells were nucleofected as previously described^57^ with slight modification using the Lonza 4D-Nucleofector core unit (Cat. No. AAF-1003B), Lonza 4D-nucleofector X unit (Cat. No. AAF-1003X), and Lonza P4 primary cell X kit L (Cat. No. V4XP-4024). Prior to nucleofection, sgRNA was resuspended overnight at 4°C in TE buffer at a concentration of 4 μg/μL. Alt-R HDR modified ssODN was resuspended overnight RT in D-PBS at a concentration of 200 pmol/μL. For each nucleofection, 4 μL of sgRNA and 2 μL of HiFi Cas9 nuclease (IDT) was incubated in a sterile cryovial for 30 minutes at RT to assemble RNPs. During the RNP incubation, KOLF2.1J-iTF cells were lifted to single cell suspension and counted as described above. 800,000 cells were pelleted at 300g for 5 minutes and resuspended in 100 μL of a nucleofection master-mix containing: 6 μL RNP assemblies, 82 μL Lonza nucleofection solution, 18 μL Lonza nucleofection supplement, and 1 μL ssODN repair template. Cells resuspended in nucleofection master-mix were transferred to a cuvette and immediately nucleofected using P4 primary nucleofection program CM113. Using a Lonza disposable Pasteur pipette, cells were then immediately transferred to a Synthemax coated 6-well plate containing 1:100 RevitaCell ROCK inhibitor and 0.5 μM Alt-R HDR Enhancer v2 (IDT, 10007910) in StemFlex media. The bottom of the cuvette was washed with extra media to collect remaining cells and plated as well. The plated, nucleofected cells were then subjected to a 48-hour 32°C / 5% CO^2^ cold shock. After 24 hours, RevitaCell and ALT-R HDR Enhancer v2 removed and replaced with fresh media. After 48 hours, cells were moved to 37°C / 5% CO^2^ and grown to 80-90% confluency.

### Generation of APOE4+/+ Monoclones

Once nucleofected cells were 80-90% confluent, cells were lifted to single cell suspension as described above. Living cells were then single cell sorted using a Fortessa flow cytometer into Synthemax coated wells of a 96-well plate containing StemFlex media and 1:100 RevitaCell. Wells were checked daily using brightfield microscopy and non-monoclonal wells were identified and excluded. Once wells reached confluency, monoclones were passaged one-to-one into Synthemax coated 24-well plates to expand further for genotyping and freezing. Cells were pelleted at 300g for 5 minutes, media was aspirated, and cells were collected for genomic DNA (gDNA) extraction using DNeasy blood & tissue kit (Cat. No. 69504). The *APOE4*^+/+^ edit site was then PCR amplified from monoclonal gDNA using the forward primer: 5’-CTGGAGGAACAACTGACCCC-3’ and the reverse primer: 5’- CTCGAACCAGCTCTTGAGGC-3’ with NEBNext High-Fidelity 2X PCR Master Mix (New England Biolabs, Cat. No. M0541S). Successful PCR amplification was confirmed by running product on 1% agarose gels. PCR products were then column purified using a DNA clean and concentrator-5 kit (Zymo, Cat. No. D4013). Purified PCR product was then submitted to GENEWIZ for sanger sequencing using the reverse PCR primer for sequencing (5’- CTCGAACCAGCTCTTGAGGC-3’). Results were aligned to the endogenous sequence using SnapGene and chromatograms were analyzed for successful ssODN mediated HDR. Of the first 12 monoclones sanger sequenced, all were successfully cut and 3 (25%) had successful biallelic HDR edits.

### KOLF2.1J-iTF iMGL Differentiation

Following previously describe methods^31,32^, KOLF2.1J-iTF iPSCs were seeded onto double coated 6-well plates (Poly-D-Lysine (Gibco, Cat. No. A3890401) coated + Matrigel (Corning, Cat. No. 356231) coated) at densities of 135K for *APOE3^+/+^* and 150K for *APOE4*^+/+^ in day 0 medium containing: Essential 8 medium (Gibco, Cat. No. A1517001), 2 μg/mL doxycycline (Takara, Cat. No. 631311), and 1X RevitaCell ROCK inhibitor. On day 2, media was aspirated and replaced with day 2 media containing: Advanced DMEM/F12, 1X GlutaMAX^TM^ Supplement (Gibco, Cat. No. 35050061), 2 μg/mL doxycycline, 100 ng/mL human IL-34 (PeproTech, Cat. No. 200-34), and 10 ng/mL human GM-CSF (PeproTech, Cat. No. 300-03). On day 4, day 2 media was aspirated and replaced with day 4 media containing: Advanced DMEM/F12, 1X GlutaMAX^TM^ Supplement, 2 μg/mL doxycycline, 100 ng/mL human IL-34, 10 ng/mL human GM-CSF, 50 ng/mL human M-CSF (PeproTech, Cat. No. 300-25), and 50 ng/mL human TGF-beta 1 (PeproTech, Cat. No. 100-21). After day 4, media was aspirated and replaced with fresh day 4 media every four days (day 8, day 12, etc.) for the remainder of the iMGL differentiation.

### Arsenic Treatment

iMGLs were differentiated as described above and on day 8 the current media was aspirated and replaced with day 4 media supplemented with either vehicle (H^2^O), .05 μM, .1 μM, .5 μM, 1 μM or 5 μM arsenic resuspended in cell culture grade water. iMGLs were then exposed to arsenic for 72 hours. Following arsenic exposure, cells were lifted using Accutase for respective assays.

### Survival Assay

iPSCs were differentiated into microglia as described above and on day 8 either vehicle (H2O) or 0.01 μM, 0.1 μM, 1 μM, 5 μM, 10 μM, 50 μM, 100 μM arsenic was added.

After a 72-hour exposure, supernatant and lifted cells were collected, combined, and counted using the Trypan blue method. Technical triplicate counts were taken and averaged per biological replicate. Student’s t-test was performed between genotypes at each concentration to determine significance.

### Phagocytic Assay

*APOE3^+/+^* and *APOE4*^+/+^ KOLF2.1J-iTF cells were seeded and differentiated as described above. On day 8, either the appropriate concentration of arsenic or vehicle (H^2^O) was added to day 8 media change. After 72 hours (day 11), .0001% GFP labelled 1 μm latex beads (Sigma, Cat. No. L1030-1ML) were added to warmed Advanced DMEM/F-12 and incubated with cells for 2 hours. After, cells were washed once with DPBS to remove excess beads and then lifted with StemPro Accutase, spun down at 300g for 5 minutes, and resuspended in 1mL Advanced DMEM/F12. Cells were put on ice and taken for fluorescent activated cell sorting (FACS) with a Fortessa flow cytometer. Tubes were vortexed prior to sorting, and 10,000 cells were counted for fluorescent quantification per replicate. Outliers were identified using ROUT method (robust regression and outlier removal), with Q = 1%. Data were analyzed using ordinary two-way ANOVA with genotype and concentration as fixed factors, followed by Bonferroni-corrected multiple comparisons.

### Immunocytochemistry

KOLF2.1J-iTF iPSCs were seeded on double coated coverslips (Electron Microscopy Sciences, Cat. No. 50-948-975) in 6-well plates and differentiated into iMGLs as described above. Once differentiated, iMGLs on coverslips were fixed in 4% paraformaldehyde for 15 minutes, washed 3x with phosphate-buffered saline (PBS) (Gibco, Cat. No. 10010023) and either were stained immediately or stored parafilmed at 4°C for up to a month in PBS. Cell membranes were permeabilized by incubating coverslips at RT in PBS with .3% Tween 20 (Sigma, Cat. No. P9416-100ML) for 5 minutes and then washed 3x with PBS for 5 minutes. Coverslips were then incubated at RT for 30 minutes in blocking buffer containing: .1g bovine serum albium (Sigma, Cat. No. 10735078001), 1 mL normal goat serum (Vector Labs, Cat. No. S-1000-20), 1.5 mL 2% Triton X-100 diluted in PBS, and 7.5 mL PBS. 1° antibodies were diluted to a desired concentration in blocking buffer and then incubated with coverslips at RT for 1 hour. Coverslips were then washed 3x with PBS and incubated at RT for 1 hour with 2° antibodies diluted 1:1000 in PBS containing .3% Tween 20. Coverslips were again washed with 3x PBS and then mounted on microscope slides using ProLong™ Diamond Antifade Mountant with DAPI (Invitrogen, Cat. No. P36966) overnight in the dark.

### Mitochondrial Assay

iPSCs were differentiated to day 8 microglia as described above and presented with either a 72-hour vehicle or arsenic exposure. After 72 hours, cells were stained with 100 nM MitoTracker Green FM (Thermo, Cat. No. M46750) and 100 nM Tetramethylrhodamine Ethyl Ester Perchlorate (TMRE) (Thermo, Cat. No. T669) at 37°C for 30 minutes in phenol red free DMEM/F12 (Thermo. Cat. No. 21041025). Cells were then washed with DPBS and stained with 1 μM MitoSox Red (Thermo, Cat. No. M36008) at 37°C for 15 minutes in HBSS^+/+^ (Thermo, Cat. No. 14025092). Cells were washed again with DPBS, lifted using Accutase, spun down at 300g for 5 minutes, supernatant aspirated, and resuspended for flow cytometry. Outliers were identified using ROUT method (robust regression and outlier removal), with Q = 1%. Data were analyzed using ordinary two-way ANOVA with genotype and concentration as fixed factors, followed by Bonferroni-corrected multiple comparisons.

### RNA Extraction, Library Preparation, and Expression Analyses

RNA extractions were performed using RNeasy Plus Mini Kit (Qiagen, Cat. No. 74136). RNA integrity was verified using an Agilent TapeStation and concentration was verified using a Qubit fluorometer. Libraries were prepared using the Tecan Ovation RNA-Seq System V2.

RNA-seq libraries were sequenced across three runs on an Illumina NextSeq 2000 to produce 75 base pair (bp) single-end reads. The FastQC software v.012.1 was used to evaluate sequencing quality^58^. Raw sequencing reads were aligned to the Hg38 genome build with STAR v.020201 and subsequently filtered for mapping quality ≥ 20 with the samtools v1.18 view function^59,60^.

The FeatureCounts command line utility was used to generate counts for NCBI RefSeq (downloaded January 2020) genes^61^. Principal component analysis was performed with the stats package in the R Statistical Programming Environment v.4.4.0 and visualized with ggplot2^62,63^. Differential expression analysis was performed with DeSeq2^64^. Genes with absolute fold-change ≥2 and FDR < 0.05 were deemed differentially expressed. The z-score standardized expression values were visualized with the R ComplexHeatmap package^65^.

Co-expression analysis was conducted to identify gene modules with similar expression patterns in the day 11 samples. Prior to analysis Combat-Seq was applied to raw counts to adjust for batch effects derived from sequencing runs^66^. Normalized counts were generated with the DeSeq2 Trimmed mean of M-values (TMM) method. Genes with an expression values < 10 counts in half of the samples were discarded. Expression values were subjected to the DeSeq2 variance stabilizing transform and the top 75% most variable genes were retained. The weighted gene co-expression network analysis (WGCNA) R package was employed with the following parameters: power=17, networkType=”signed’, corType=”bicor”, minModuleSize=30, mergeCloseModules=0.25^67^. A post-analysis filter was applied to maintain genes with module connectivity (kMe) > 0.6. Differential expression analyses were further analyzed using QIAGEN Ingenuity Pathway Analysis (QIAGEN IPA). Differential expression analysis data were analyzed and the networks and functional analyses were generated through the use of QIAGEN IPA (QIAGEN Inc., https://digitalinsights.qiagen.com/IPA) using algorithms developed for use in QIAGEN IPA^68^ under gene filtering parameters of absolute fold-change ≥2 and P^adj^ < 0.05. Co-expression gene modules also were analyzed using QIAGEN IPA without gene filtering parameters.

### Whole Genome Sequencing

Whole genome sequencing was performed on paired-end libraries with 150 bp read length. Raw reads were trimmed with TrimGalore! and aligned to the Hg38 genome build with bwa mem^69,70^. Duplicate reads were tagged with the Genome Analysis Toolkit (GATK) MarkDuplicates function and filtered for mapping quality >20^71^. Read-group tags were added to alignment files with the GATK AddOrReplaceReadGroups function. GATK’s HaplotypeCaller was used to identify single-nucleotide polymorphisms (SNPs) and small insertions and deletions (indels) in variant call format (VCF). The following hard filters were applied to the raw variants: QD < 2.0, SOR > 3.0, FS > 60, MQ > 40.0, MQRankSum < −12.5, ReadPosRankSum < −8.0, DP <10 and FS < 60.0 (SNPs) or < 200.0 (indels). The bcftools v.1.18 command line utility was used to normalize vcfs and PASS variants were extracted^72^. Unique and intersecting variants were visualized as a venn diagram using the R Venndir package^73^. Functional consequences of mutations were predicted with ANNOVAR v.2020Jun08^74^.

### Volcano plot generation

Volcano plots were generated for the *APOE4*^+/+^ versus *APOE3^+/+^* differential expression datasets using a custom python workflow. The same analysis and visualization criteria were applied to all datasets. Genes were first filtered to remove low-count features using a baseMean threshold of ≥ 10. Genes with missing or non-finite values in the log2FoldChange, padj, or Fold_Change fields were excluded prior to plotting. For visualization, the x-axis was defined as log^2^(Fold-Change) and the y-axis as −log^10^(p^adj^). Statistical significance was defined as p^adj^ ≤ 0.05 together with an absolute linear fold change ≥ 2 or ≤ −2, using the Fold_Change values within the datasets.

## Authorship contribution statement

EG, AJM, and AMB conceived of the study. AJM performed experiments. AMB and AJM analyzed sequencing data. AJM, EG, and AMB wrote the manuscript with input from ET, MC, and SD.

## Acknowledgments

The authors would like to thank Carl Bortner for help with flow cytometry, the staff in the NIEHS Fluorescence Microscopy and Imaging Center for their help with imaging, and Molly Cook and the staff at the Genomics Core for help with library prep and sequencing.

## Data availability

Raw whole genome and RNA-sequencing data are available from the NCBI Sequence Read Archive (SRA) database under BioProject PRJNA1461146.

