## Supplementary Figures 1-7 for "Alzheimer’s Disease Risk Allele *APOE4* Interacts with Arsenic Exposure to Drive Microglial Dysfunction"

**Supplemental Figure Legends:**

**Supplementary Figure 1:** (A) Sanger sequencing of APOE3+/+ control and APOE4+/+ monoclones aligned to the endogenous APOE4 locus with representative chromatogram shown. (B) Ven diagram depicting shared and novel mismatches to hg38 from whole genome sequencing of two APOE4+/+ and the parental APOE3+/+ cell line. (C) Graph depicting the type and proportion of mutations detected from whole genome sequencing. (D) Heatmap of gene expression changes between APOE3+/+ and APOE4+/+ iTFMs on differentiation day 8. (E) Volcano plot of differentially expressed genes in APOE4+/+ compared to APOE3+/+.

**Supplementary Figure 2:** Volcano plots of 72 hr arsenite treated APOE3+/+ iTFM compared to vehicle treated APOE3+/+ iTFM.

**Supplementary Figure 3:** Volcano plots of 72 hr arsenite treated APOE4+/+ iTFM compared to vehicle treated APOE4+/+ iTFM.

**Supplementary Figure 4:** Volcano plots of 72 hr vehicle and arsenite treated APOE4+/+ iTFM compared to the respective APOE3+/+ control iTFM.

**Supplementary Figure 5:** (A-F) Ingenuity pathway analysis of APOE4+/+ differentially expressed genes (absolute fold-change ≥2, padj < 0.05) when treated with increasing concentrations of arsenite for 72 hrs (top to bottom). Shown are top enriched pathways (left), top predicted upstream regulators (middle), and top predicted cellular functions (right).

**Supplementary Figure 6:** PCA of weighted gene co-expression network analysis for APOE3+/+ (blue) and APOE4+/+ (yellow). Each dot represents an independent biological sample responding to increasing concentration of arsenite (light to dark).

**Supplementary Figure 7:** Mean fluorescent intensity of (C-D) APOE3+/+ (purple) and APOE4+/+ (teal) iTFM. Statistical significance assessed by two-way ANOVA followed by Bonferroni post hoc testing. *, **, and *** correspond to p values <0.05, 0.005, 0.0001 respectively.

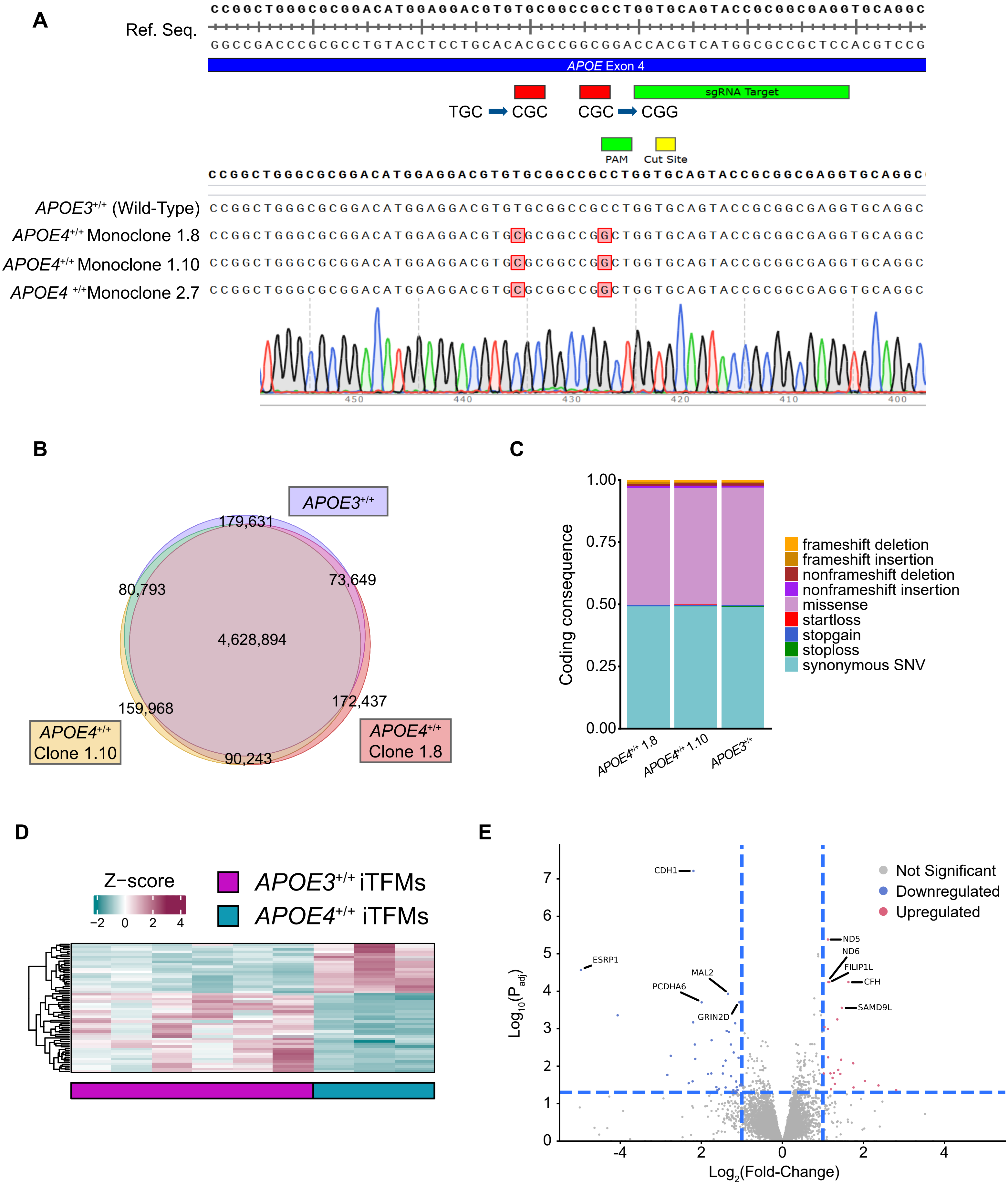
A
Ref. Seq.
CCGGCTGGGCGCGGACATGGAGGACGTGTGCGGCCGCCTGGTGCAGTACCGCGGCGAGGTGCAGGC
GGCCGACCCGCGCCTGTACCTCCTGCACACGCCGGCGGACCACGTCATGGCGCCGCTCCACGTCCG
APOE Exon 4
sgRNA Target
TGC ➡ CGC
CGC ➡ CGG
PAM
Cut Site
CCGGCTGGGCGCGGACATGGAGGACGTGTGCGGCCGCCTGGTGCAGTACCGCGGCGAGGTGCAGGC
APOE3+/+ (Wild-Type)
APOE4+/+ Monoclone 1.8
APOE4+/+ Monoclone 1.10
APOE4 +/+ Monoclone 2.7
450
440
430
420
410
400
B
APOE3+/+
179,631
80,793
73,649
4,628,894
159,968
172,437
90,243
APOE4+/+ Clone 1.10
APOE4+/+ Clone 1.8
C
Coding consequence
1.00
0.75
0.50
0.25
0.00
APOE4+/+ 1.8
APOE4+/+ 1.10
APOE3+/+
frameshift deletion
frameshift insertion
nonframeshift deletion
nonframeshift insertion
missense
startloss
stopgain
stoploss
synonymous SNV
D
Z−score
−2 0 2 4
APOE3+/+ iTFMs
APOE4+/+ iTFMs
E
Not Significant
Downregulated
Upregulated
CDH1
ESRP1
MAL2
PCDHA6
GRIN2D
ND5
ND6
FILIP1L
CFH
SAMD9L
Log10(Padj)
Log2(Fold-Change)

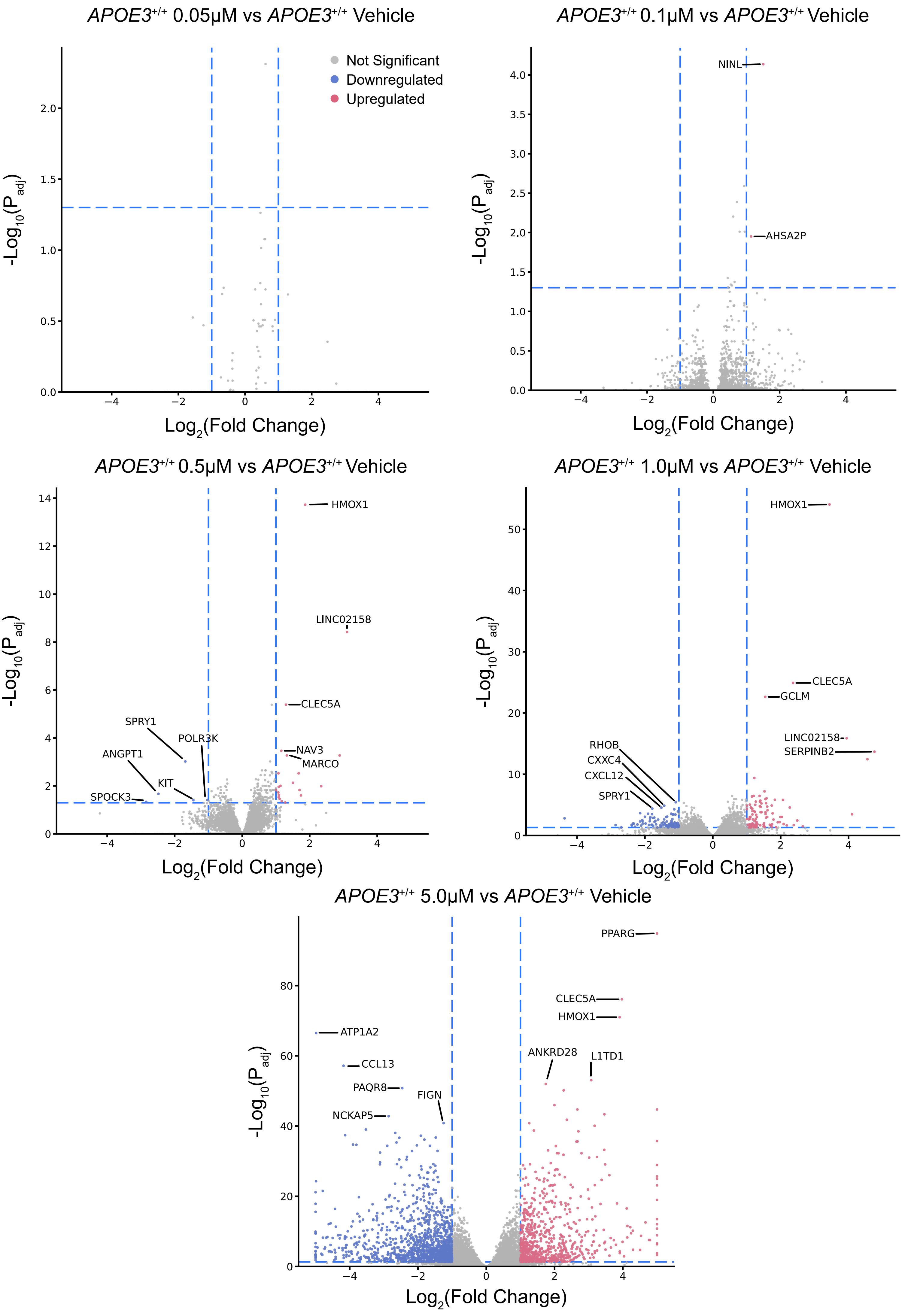

APOE3+/+ 0.05μM vs APOE3+/+ Vehicle
Not Significant
Downregulated
Upregulated
-Log10(Padj)
Log2(Fold Change)
APOE3+/+ 0.1μM vs APOE3+/+ Vehicle
NINL
AHSA2P
-Log10(Padj)
Log2(Fold Change)
APOE3+/+ 0.5μM vs APOE3+/+ Vehicle
HMOX1
LINC02158
CLEC5A
SPRY1
POLR3K
ANGPT1
NAV3
MARCO
KIT
SPOCK3
-Log10(Padj)
Log2(Fold Change)
APOE3+/+ 1.0μM vs APOE3+/+ Vehicle
HMOX1
CLEC5A
GCLM
LINC02158
SERPINB2
RHOB
CXXC4
CXCL12
SPRY1
-Log10(Padj)
Log2(Fold Change)
APOE3+/+ 5.0μM vs APOE3+/+ Vehicle
PPARG
CLEC5A
HMOX1
ATP1A2
CCL13
PAQR8
FIGN
NCKAP5
ANKRD28
L1TD1
-Log10(Padj)
Log2(Fold Change)

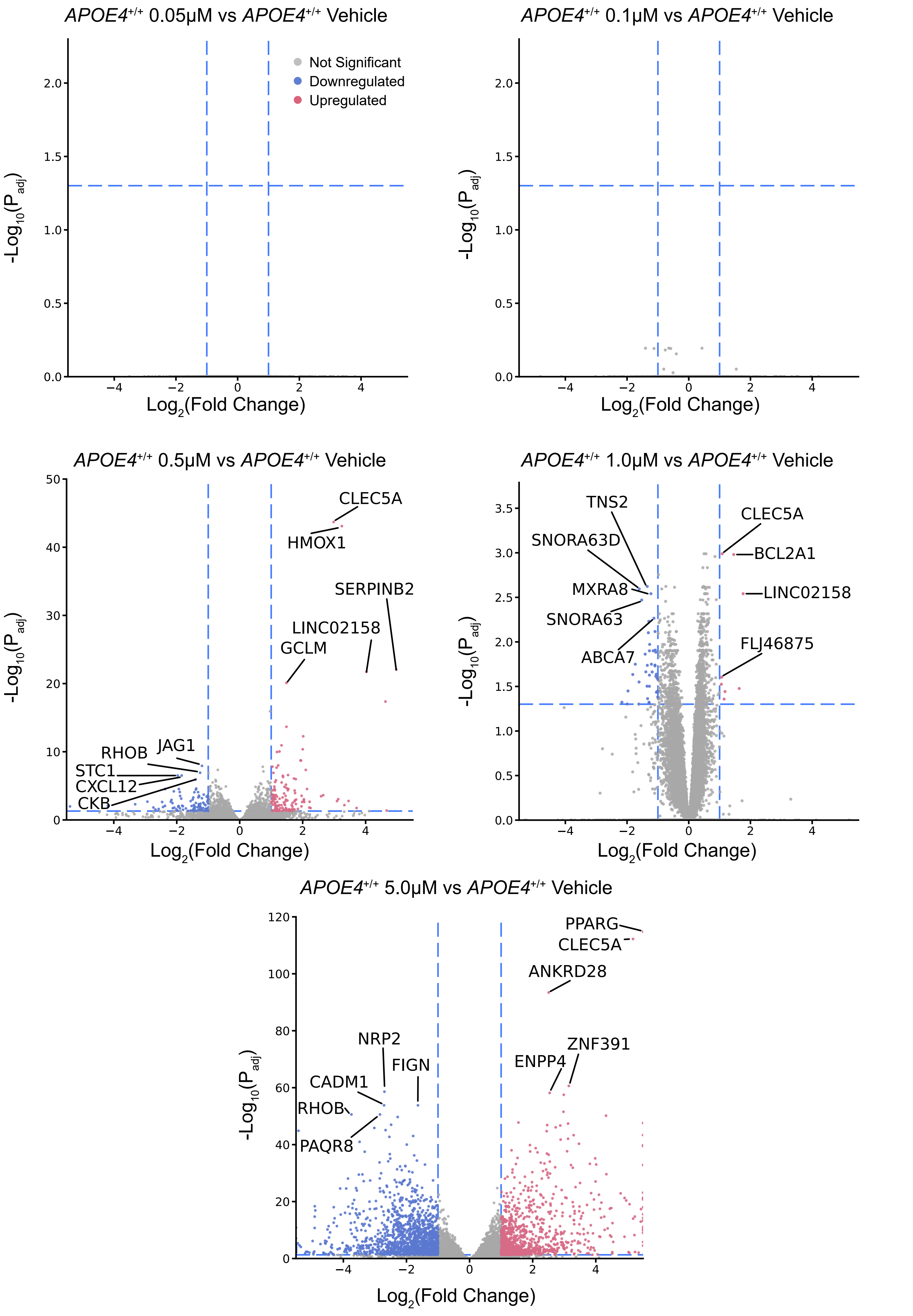
APOE4+/+ 0.05μM vs APOE4+/+ Vehicle
Not Significant
Downregulated
Upregulated
-Log10(Padj)
Log2(Fold Change)
APOE4+/+ 0.1μM vs APOE4+/+ Vehicle
APOE4+/+ 0.5μM vs APOE4+/+ Vehicle
CLEC5A
HMOX1
SERPINB2
LINC02158
GCLM
RHOB
JAG1
STC1
CXCL12
CKB
APOE4+/+ 1.0μM vs APOE4+/+ Vehicle
TNS2
SNORA63D
MXRA8
SNORA63
ABCA7
CLEC5A
BCL2A1
LINC02158
FLJ46875
APOE4+/+ 5.0μM vs APOE4+/+ Vehicle
PPARG
CLEC5A
ANKRD28
NRP2
FIGN
CADM1
RHOB
PAQR8
ENPP4
ZNF391

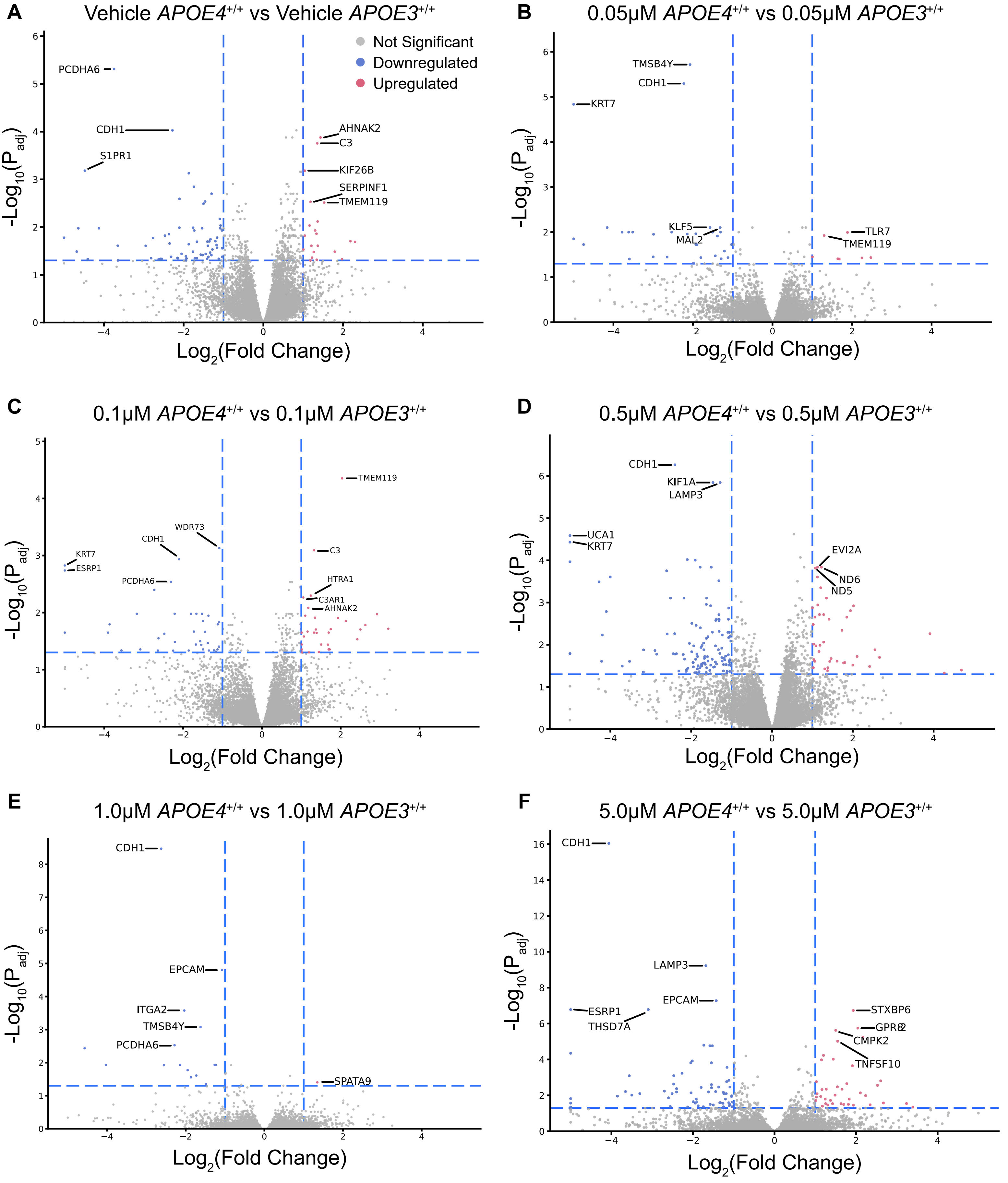
A
Vehicle APOE4+/+ vs Vehicle APOE3+/+
Not Significant
Downregulated
Upregulated
PCDHA6
CDH1
S1PR1
AHNAK2
C3
KIF26B
SERPINF1
TMEM119
-Log10(Padj)
Log2(Fold Change)
B
0.05μM APOE4+/+ vs 0.05μM APOE3+/+
TMSB4Y
CDH1
KRT7
KLF5
MAL2
TLR7
TMEM119
-Log10(Padj)
Log2(Fold Change)
C
0.1μM APOE4+/+ vs 0.1μM APOE3+/+
TMEM119
WDR73
CDH1
KRT7
ESRP1
PCDHA6
C3
HTRA1
C3AR1
AHNAK2
-Log10(Padj)
Log2(Fold Change)
D
0.5μM APOE4+/+ vs 0.5μM APOE3+/+
CDH1
KIF1A
LAMP3
UCA1
KRT7
EVI2A
ND6
ND5
-Log10(Padj)
Log2(Fold Change)
E
1.0μM APOE4+/+ vs 1.0μM APOE3+/+
CDH1
EPCAM
ITGA2
TMSB4Y
PCDHA6
SPATA9
-Log10(Padj)
Log2(Fold Change)
F
5.0μM APOE4+/+ vs 5.0μM APOE3+/+
CDH1
LAMP3
EPCAM
ESRP1
THSD7A
STXBP6
GPR82
CMPK2
TNFSF10
-Log10(Padj)
Log2(Fold Change)

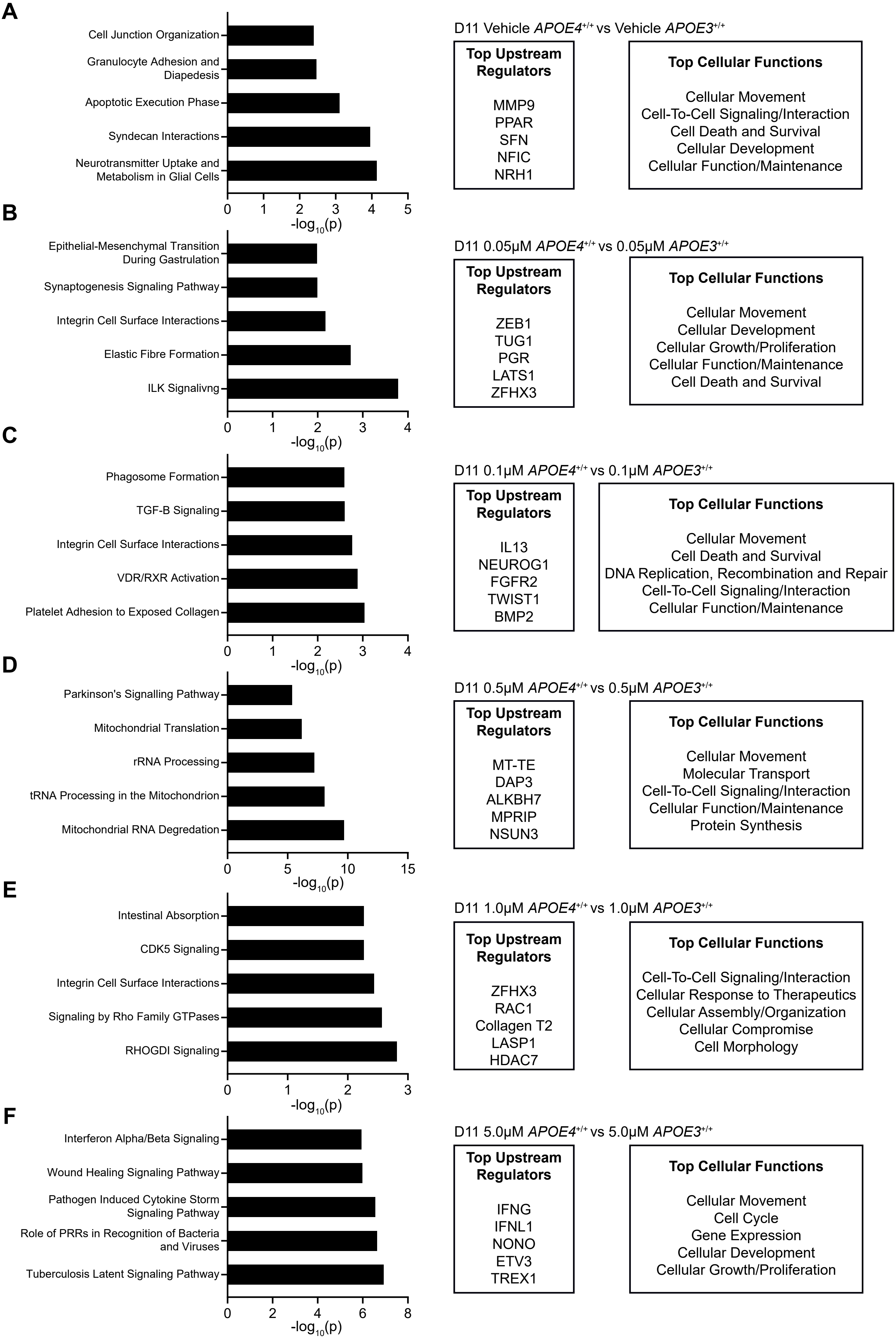
A
Cell Junction Organization
Granulocyte Adhesion and Diapedesis
Apoptotic Execution Phase
Syndecan Interactions
Neurotransmitter Uptake and Metabolism in Glial Cells
0 1 2 3 4 5
-log10(p)
D11 Vehicle APOE4+/+ vs Vehicle APOE3+/+
Top Upstream Regulators
MMP9
PPAR
SFN
NFIC
NRH1
Top Cellular Functions
Cellular Movement
Cell-To-Cell Signaling/Interaction
Cell Death and Survival
Cellular Development
Cellular Function/Maintenance
B
Epithelial-Mesenchymal Transition During Gastrulation
Synaptogenesis Signaling Pathway
Integrin Cell Surface Interactions
Elastic Fibre Formation
ILK Signalivng
0 1 2 3 4
-log10(p)
D11 0.05μM APOE4+/+ vs 0.05μM APOE3+/+
Top Upstream Regulators
ZEB1
TUG1
PGR
LATS1
ZFHX3
Top Cellular Functions
Cellular Movement
Cellular Development
Cellular Growth/Proliferation
Cellular Function/Maintenance
Cell Death and Survival
C
Phagosome Formation
TGF-B Signaling
Integrin Cell Surface Interactions
VDR/RXR Activation
Platelet Adhesion to Exposed Collagen
0 1 2 3 4
-log10(p)
D11 0.1μM APOE4+/+ vs 0.1μM APOE3+/+
Top Upstream Regulators
IL13
NEUROG1
FGFR2
TWIST1
BMP2
Top Cellular Functions
Cellular Movement
Cell Death and Survival
DNA Replication, Recombination and Repair
Cell-To-Cell Signaling/Interaction
Cellular Function/Maintenance
D
Parkinson's Signalling Pathway
Mitochondrial Translation
rRNA Processing
tRNA Processing in the Mitochondrion
Mitochondrial RNA Degredation
0 5 10 15
-log10(p)
D11 0.5μM APOE4+/+ vs 0.5μM APOE3+/+
Top Upstream Regulators
MT-TE
DAP3
ALKBH7
MPRIP
NSUN3
Top Cellular Functions
Cellular Movement
Molecular Transport
Cell-To-Cell Signaling/Interaction
Cellular Function/Maintenance
Protein Synthesis
E
Intestinal Absorption
CDK5 Signaling
Integrin Cell Surface Interactions
Signaling by Rho Family GTPases
RHOGDI Signaling
0 1 2 3
-log10(p)
D11 1.0μM APOE4+/+ vs 1.0μM APOE3+/+
Top Upstream Regulators
ZFHX3
RAC1
Collagen T2
LASP1
HDAC7
Top Cellular Functions
Cell-To-Cell Signaling/Interaction
Cellular Response to Therapeutics
Cellular Assembly/Organization
Cellular Compromise
Cell Morphology
F
Interferon Alpha/Beta Signaling
Wound Healing Signaling Pathway
Pathogen Induced Cytokine Storm Signaling Pathway
Role of PRRs in Recognition of Bacteria and Viruses
Tuberculosis Latent Signaling Pathway
0 2 4 6 8
-log10(p)
D11 5.0μM APOE4+/+ vs 5.0μM APOE3+/+
Top Upstream Regulators
IFNG
IFNL1
NONO
ETV3
TREX1
Top Cellular Functions
Cellular Movement
Cell Cycle
Gene Expression
Cellular Development
Cellular Growth/Proliferation

A

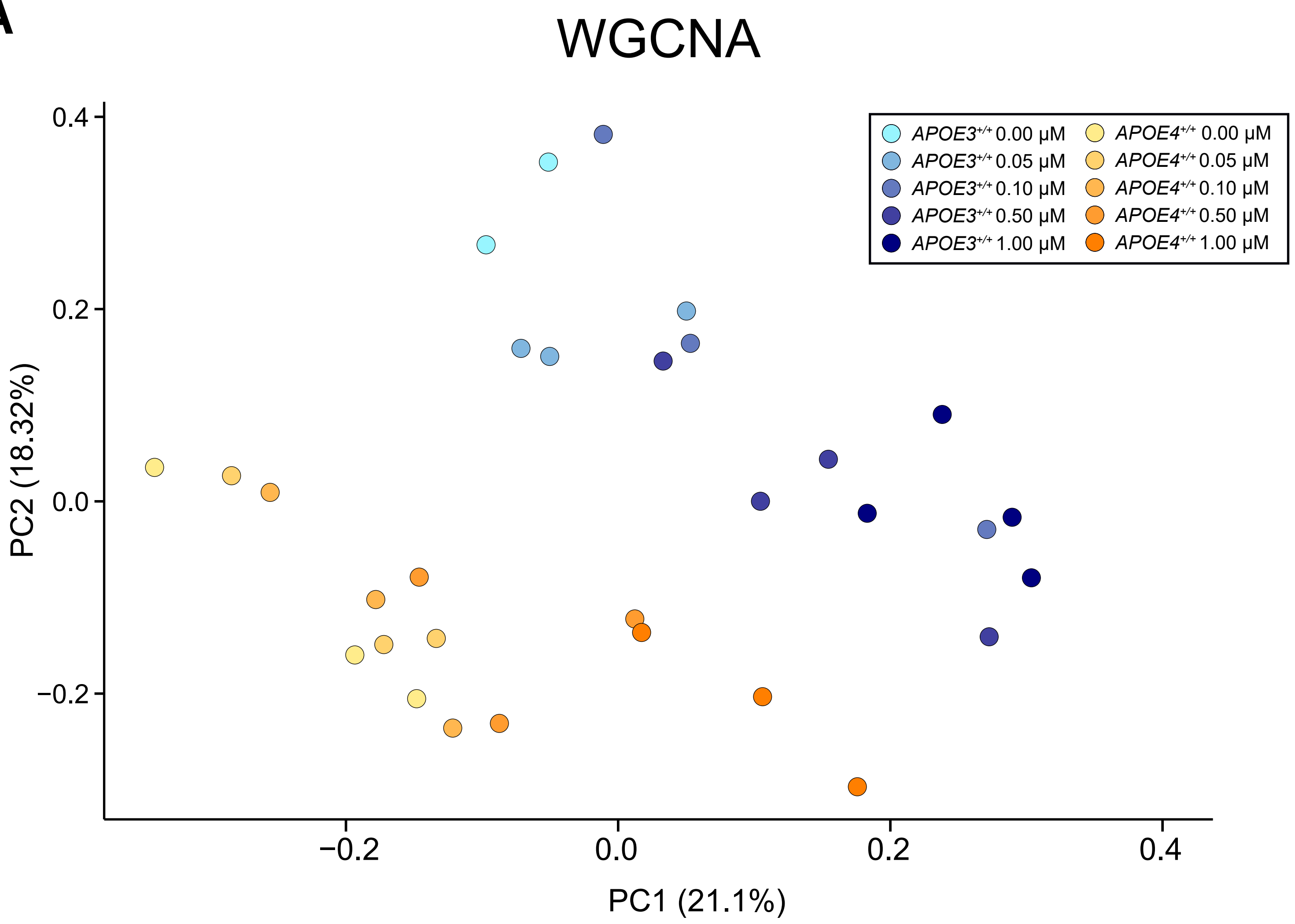

WGCNA
PC2 (18.32%)
PC1 (21.1%)
0.4
0.2
0.0
−0.2
−0.2
0.0
0.2
0.4
APOE3+/+ 0.00 μM
APOE3+/+ 0.05 μM
APOE3+/+ 0.10 μM
APOE3+/+ 0.50 μM
APOE3+/+ 1.00 μM
APOE4+/+ 0.00 μM
APOE4+/+ 0.05 μM
APOE4+/+ 0.10 μM
APOE4+/+ 0.50 μM
APOE4+/+ 1.00 μM

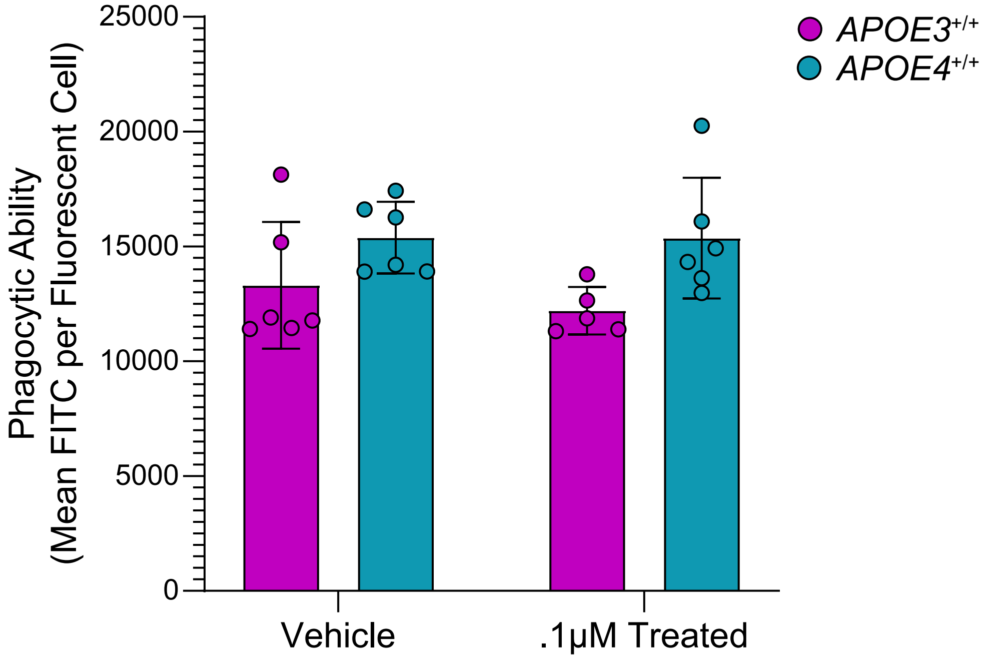

Phagocytic Ability
(Mean FITC per Fluorescent Cell)
25000
20000
15000
10000
5000
0
Vehicle
.1μM Treated
APOE3+/+
APOE4+/+
