## Supplementary Figure 8 for "Alzheimer’s Disease Risk Allele *APOE4* Interacts with Arsenic Exposure to Drive Microglial Dysfunction"

**Supplementary Figure 8:** Heatmaps of assigned WGCNA modules showing relative expression changes across arsenite concentration. *APOE3*$^{+/+}$ in pink, *APOE4*$^{+/+}$ in blue.

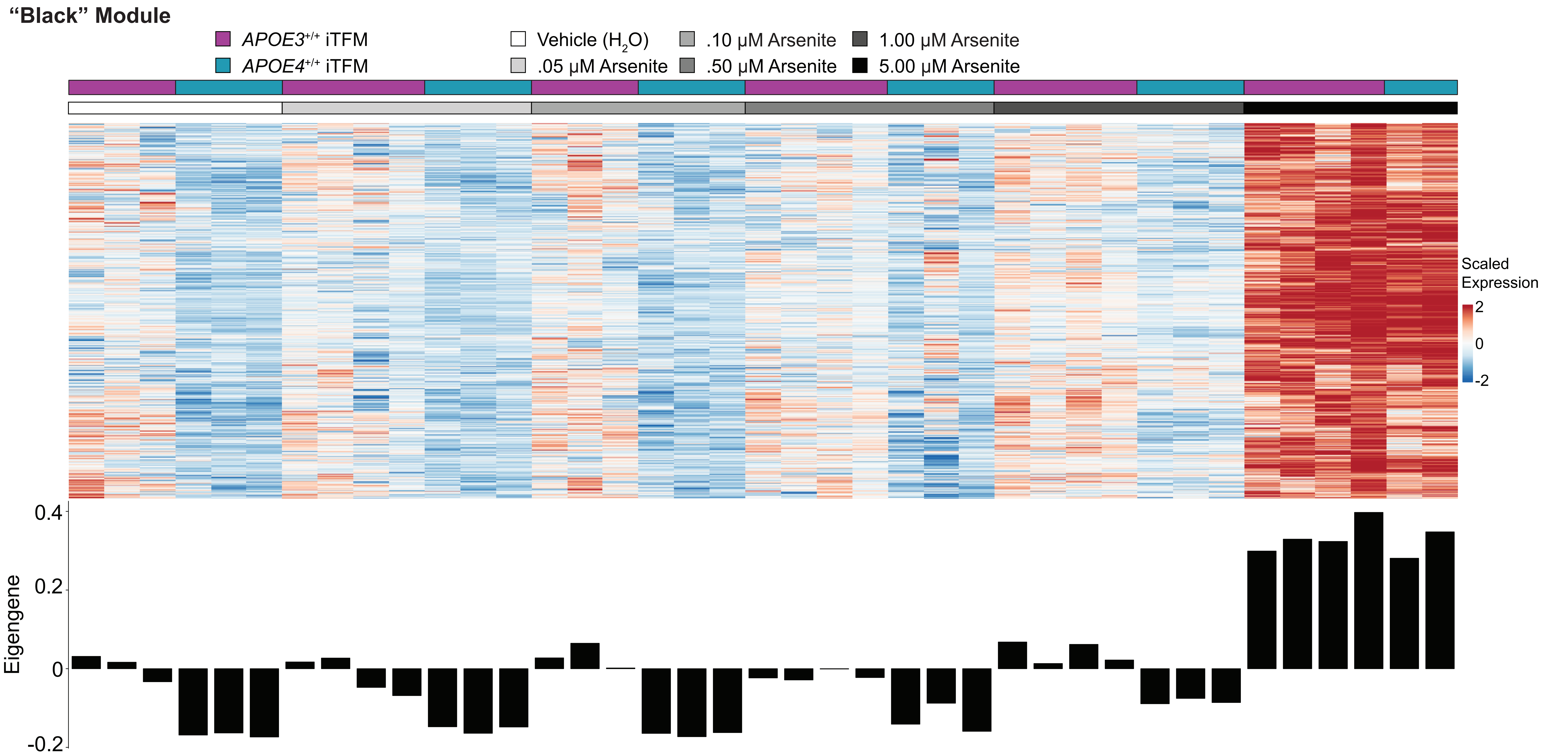

"Black" Module
APOE3+/+ iTFM
APOE4+/+ iTFM
Vehicle (H2O)
.05 μM Arsenite
.10 μM Arsenite
.50 μM Arsenite
1.00 μM Arsenite
5.00 μM Arsenite
Scaled Expression
2
0
-2
Eigengene
0.4
0.2
0
-0.2

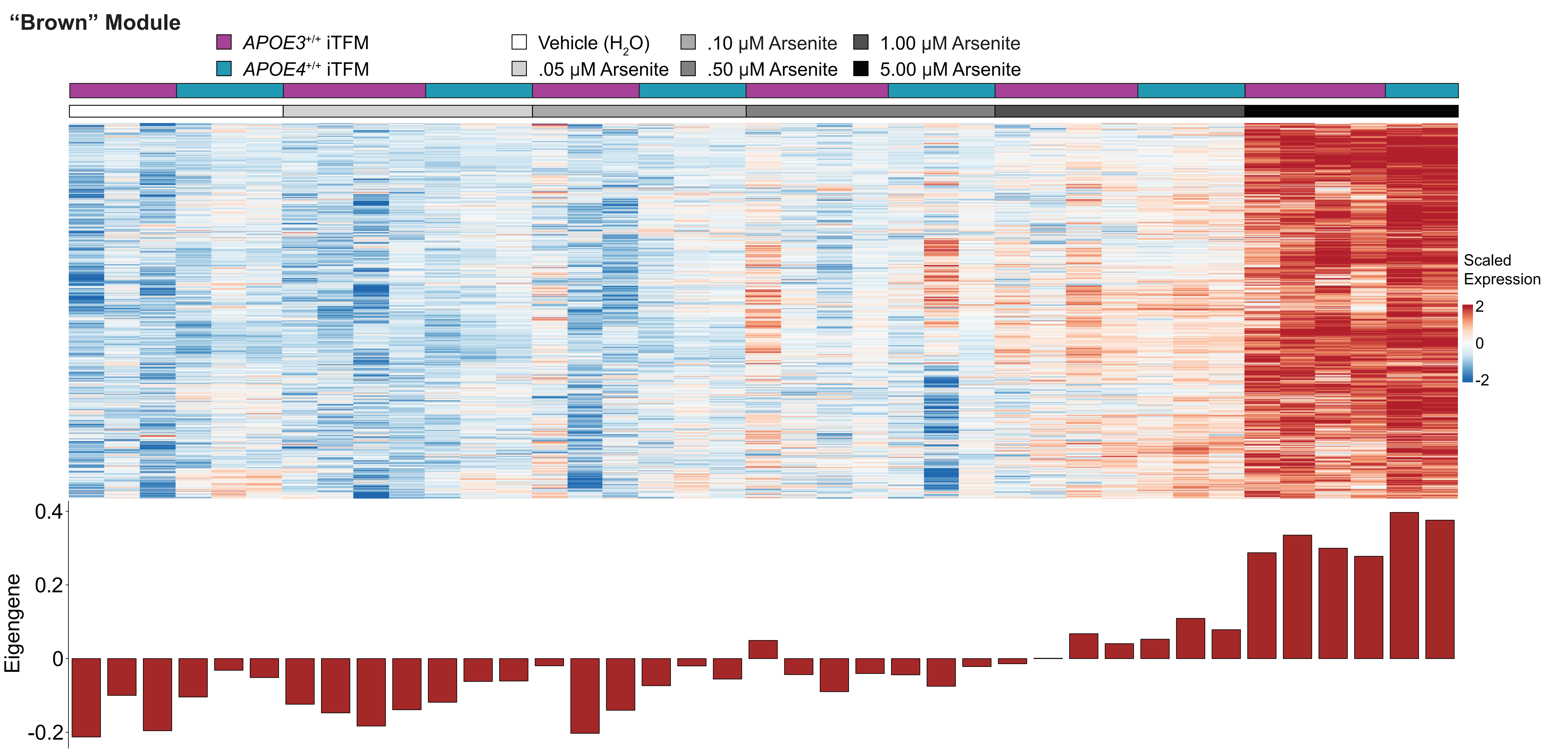

"Brown" Module
APOE3+/+ iTFM
APOE4+/+ iTFM
Vehicle (H2O)
.05 μM Arsenite
.10 μM Arsenite
.50 μM Arsenite
1.00 μM Arsenite
5.00 μM Arsenite
Scaled Expression
2
0
-2
Eigengene
0.4
0.2
0
-0.2

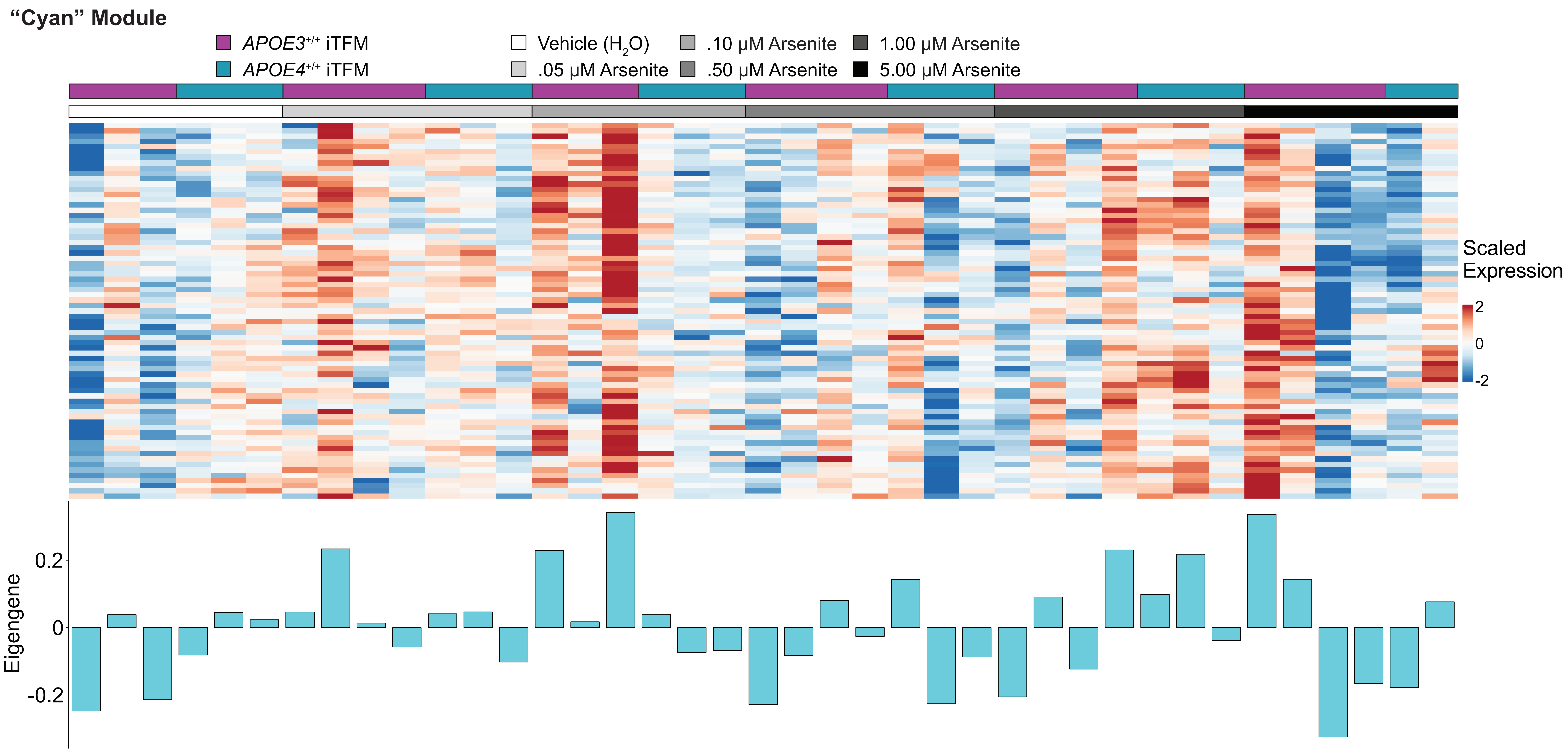
"Cyan" Module
APOE3+/+ iTFM
APOE4+/+ iTFM
Vehicle (H2O)
.05 μM Arsenite
.10 μM Arsenite
.50 μM Arsenite
1.00 μM Arsenite
5.00 μM Arsenite
Scaled Expression
2
0
-2
Eigengene
0.2
0
-0.2

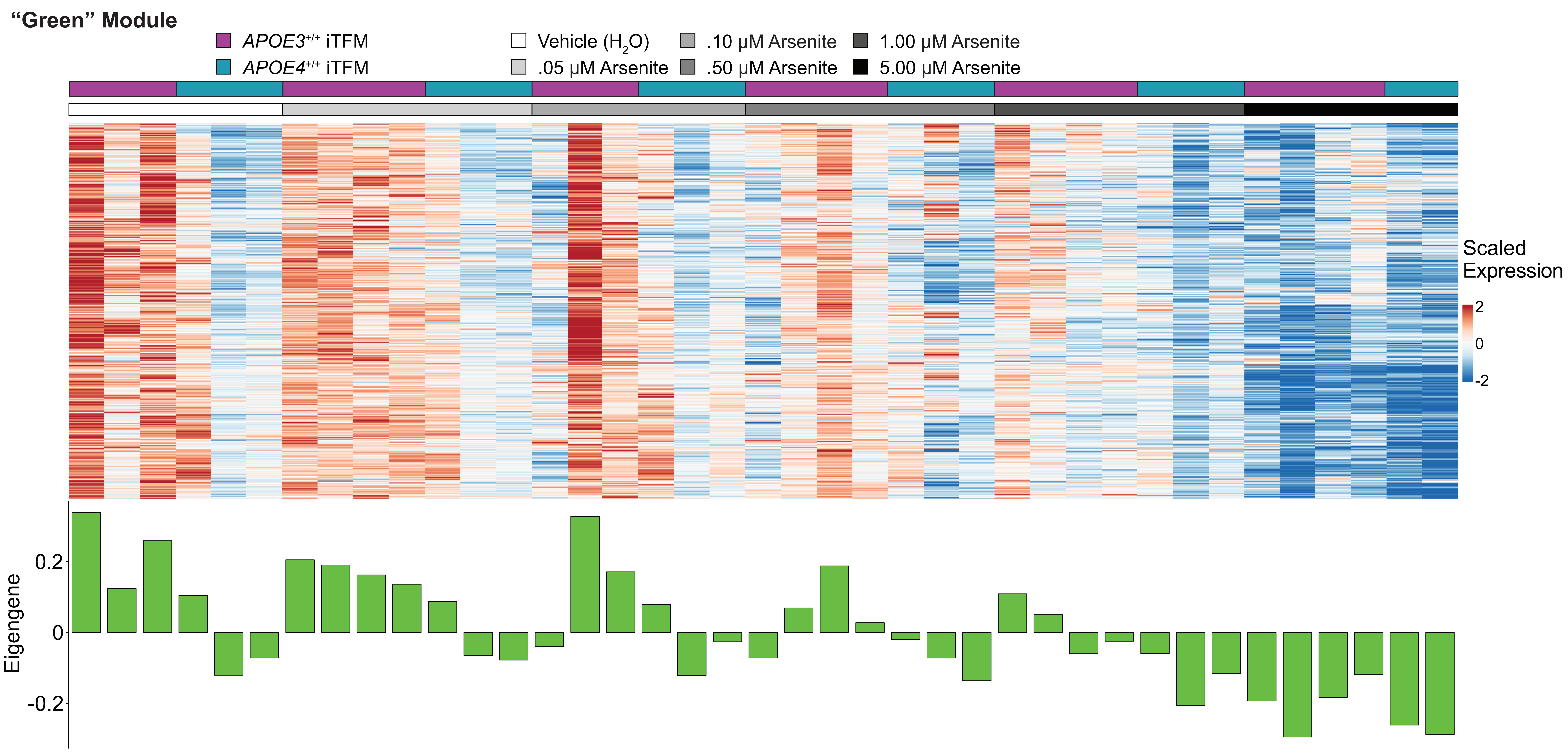
"Green" Module
APOE3+/+ iTFM
APOE4+/+ iTFM
Vehicle (H2O)
.05 μM Arsenite
.10 μM Arsenite
.50 μM Arsenite
1.00 μM Arsenite
5.00 μM Arsenite
Scaled Expression
2
0
-2
Eigengene
0.2
0
-0.2

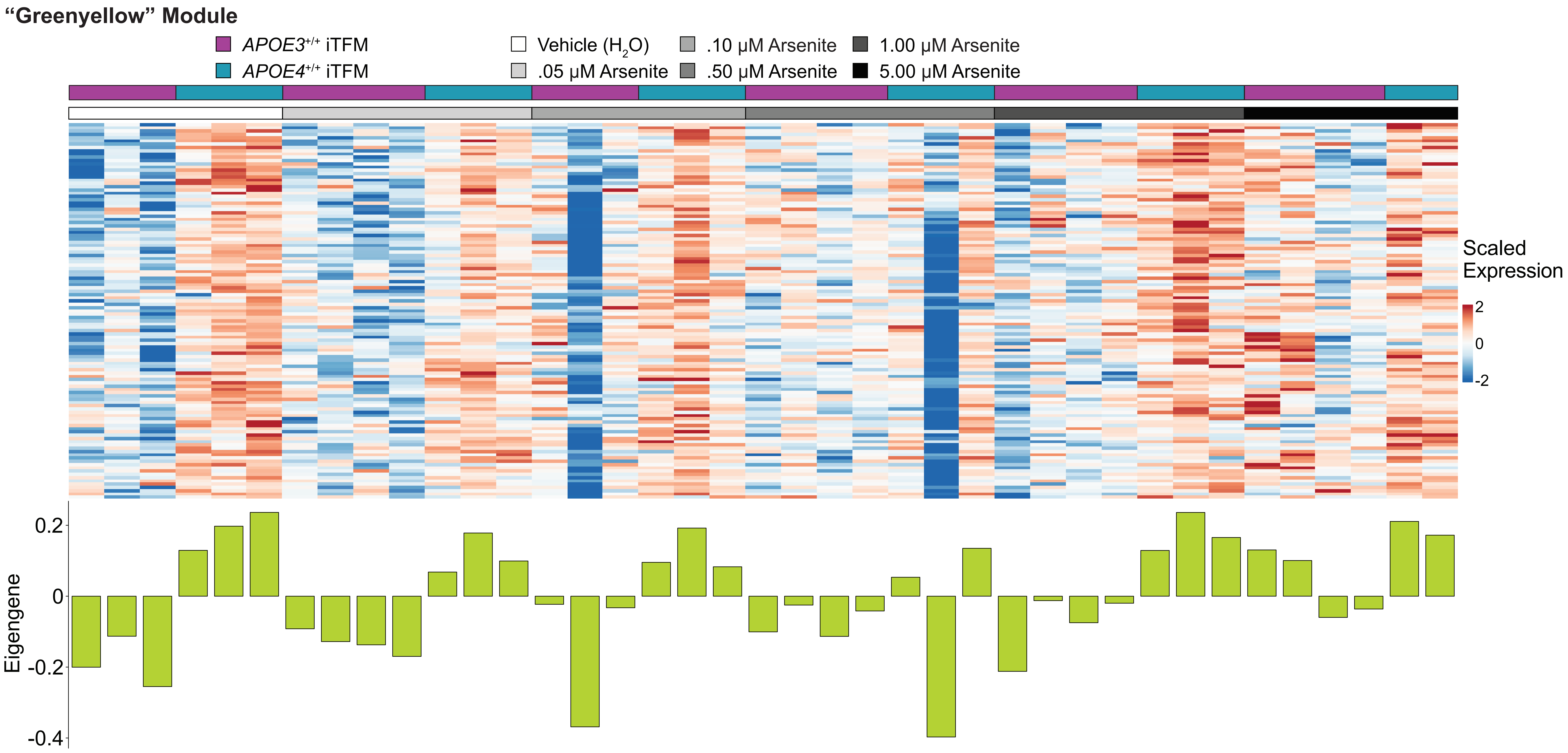
"Greenyellow" Module
APOE3+/+ iTFM
APOE4+/+ iTFM
Vehicle (H2O)
.05 μM Arsenite
.10 μM Arsenite
.50 μM Arsenite
1.00 μM Arsenite
5.00 μM Arsenite
Scaled Expression
2
0
-2
Eigengene
0.2
0
-0.2
-0.4

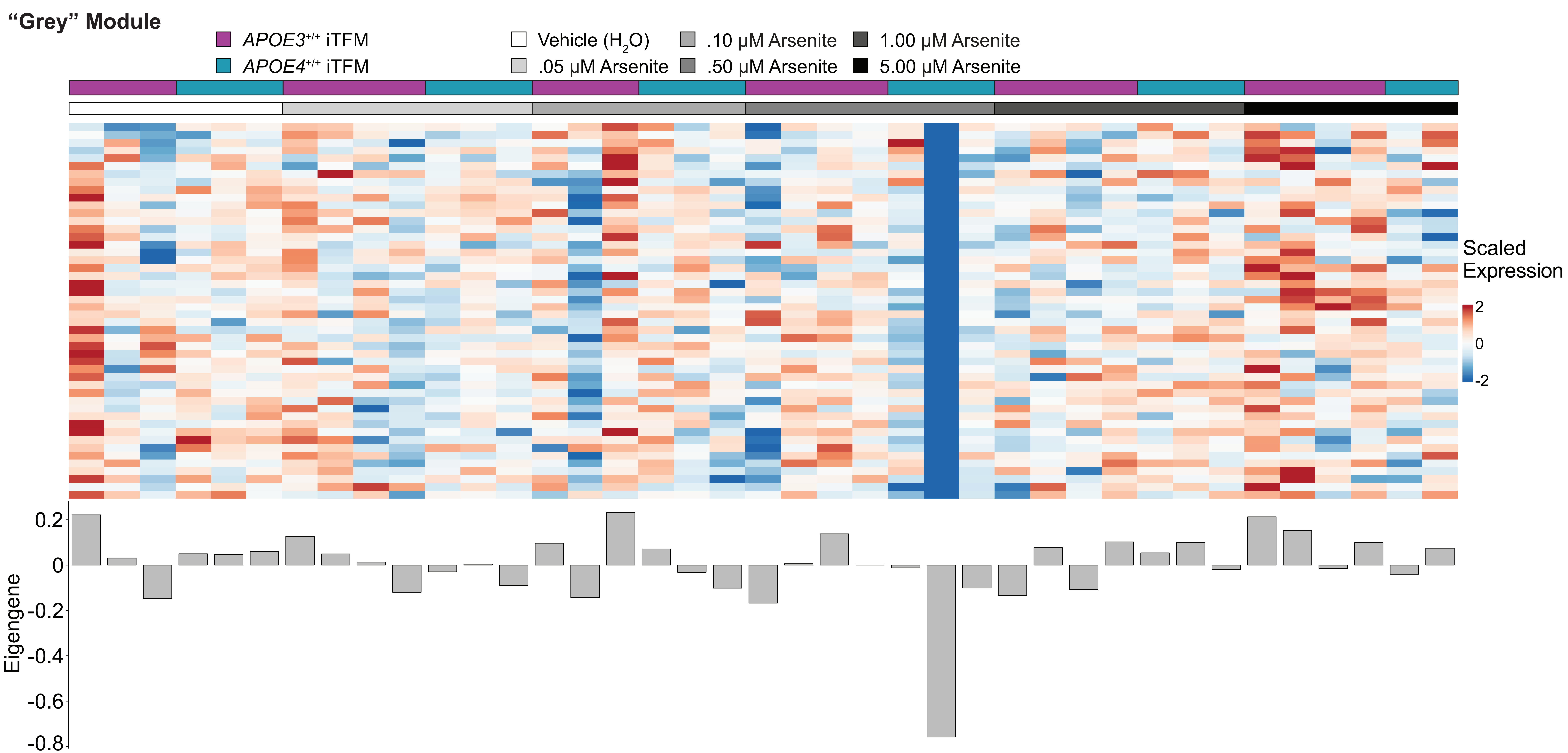

"Grey" Module
APOE3+/+ iTFM
APOE4+/+ iTFM
Vehicle (H2O)
.05 μM Arsenite
.10 μM Arsenite
.50 μM Arsenite
1.00 μM Arsenite
5.00 μM Arsenite
Scaled Expression
2
0
-2
Eigengene
0.2
0
-0.2
-0.4
-0.6
-0.8

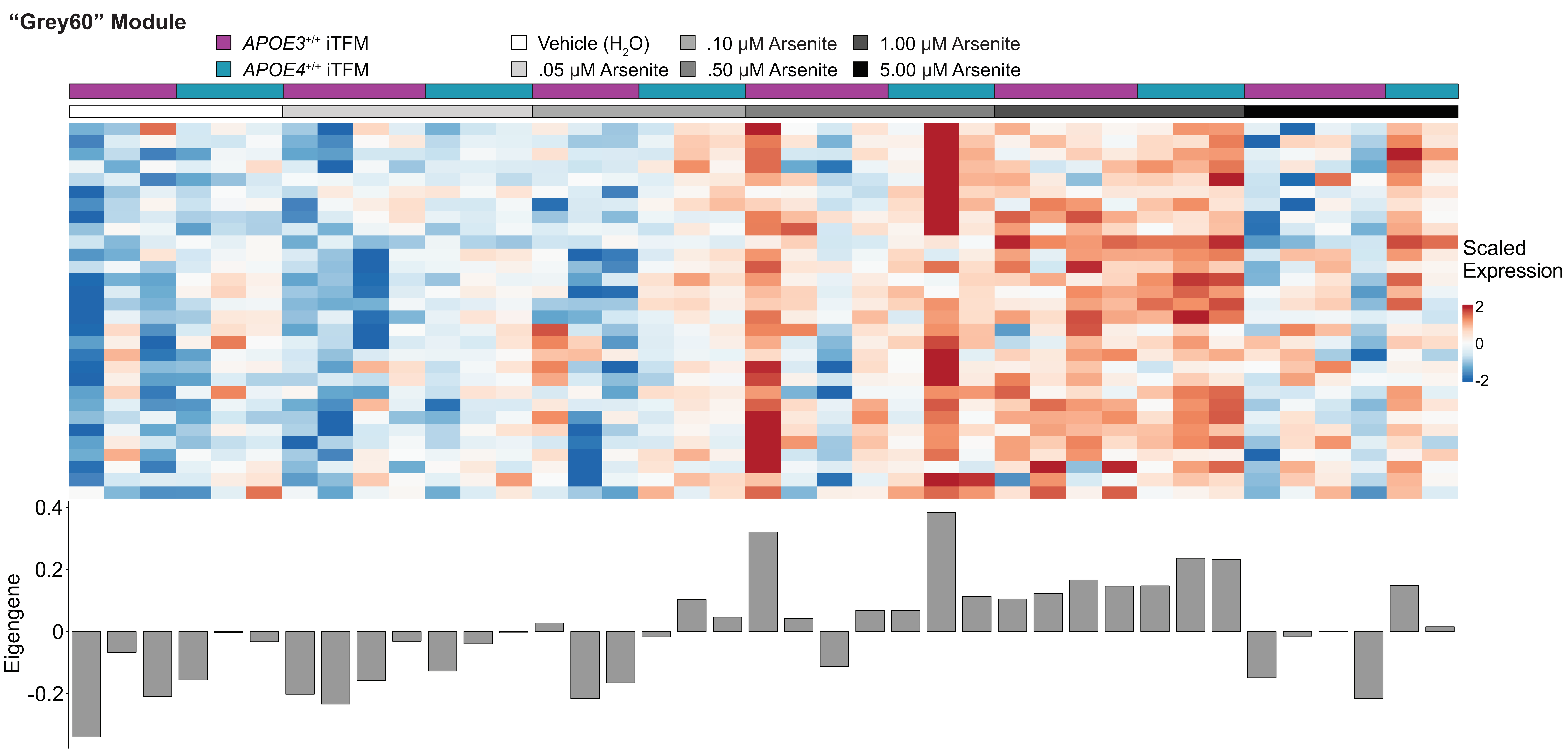
"Grey60" Module
APOE3+/+ iTFM
APOE4+/+ iTFM
Vehicle (H2O)
.05 μM Arsenite
.10 μM Arsenite
.50 μM Arsenite
1.00 μM Arsenite
5.00 μM Arsenite
Scaled Expression
2
0
-2
Eigengene
0.4
0.2
0
-0.2

### "LightCyan" Module

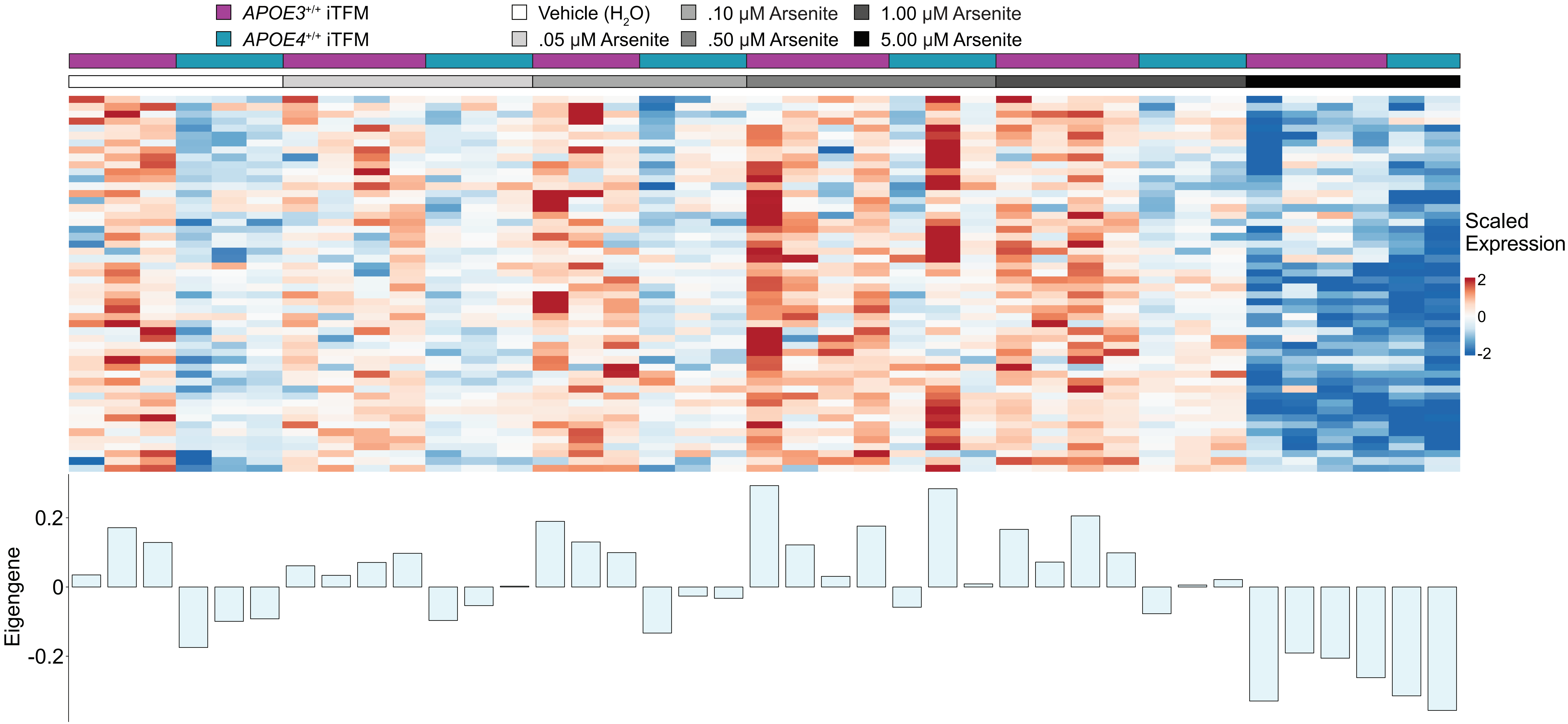

“Magenta” Module

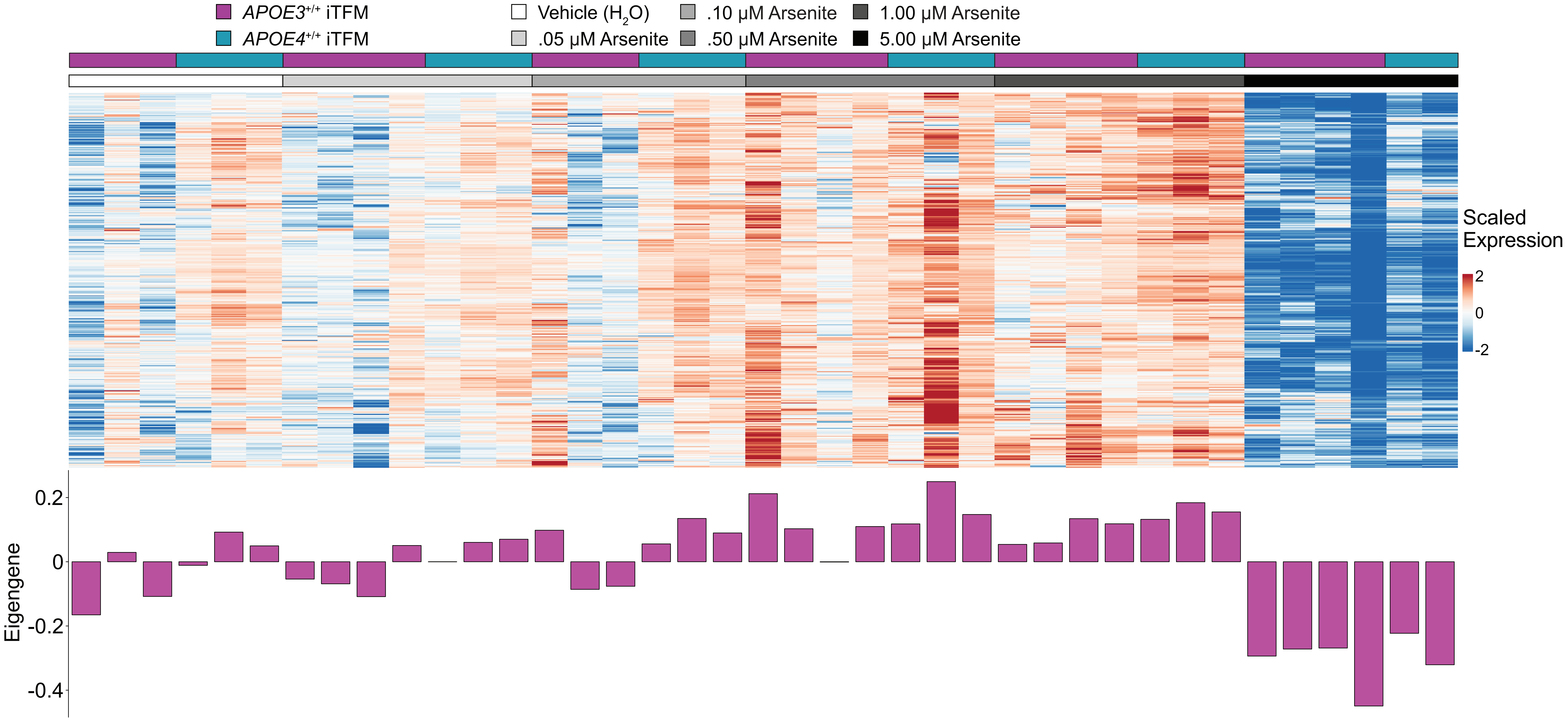

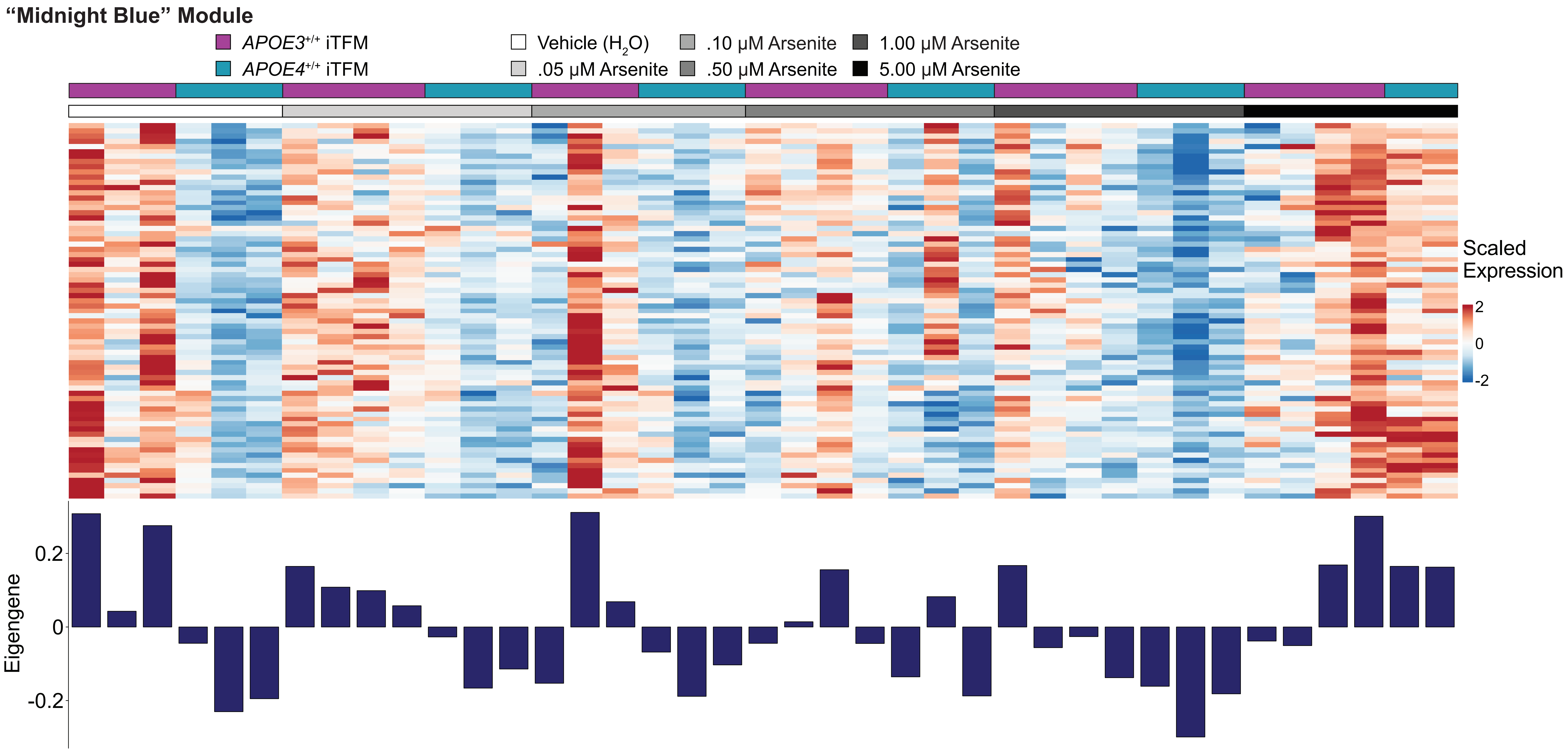
"Midnight Blue" Module
APOE3+/+ iTFM
APOE4+/+ iTFM
Vehicle (H2O)
.05 μM Arsenite
.10 μM Arsenite
.50 μM Arsenite
1.00 μM Arsenite
5.00 μM Arsenite
Scaled Expression
2
0
-2
Eigengene
0.2
0
-0.2

"Purple" Module

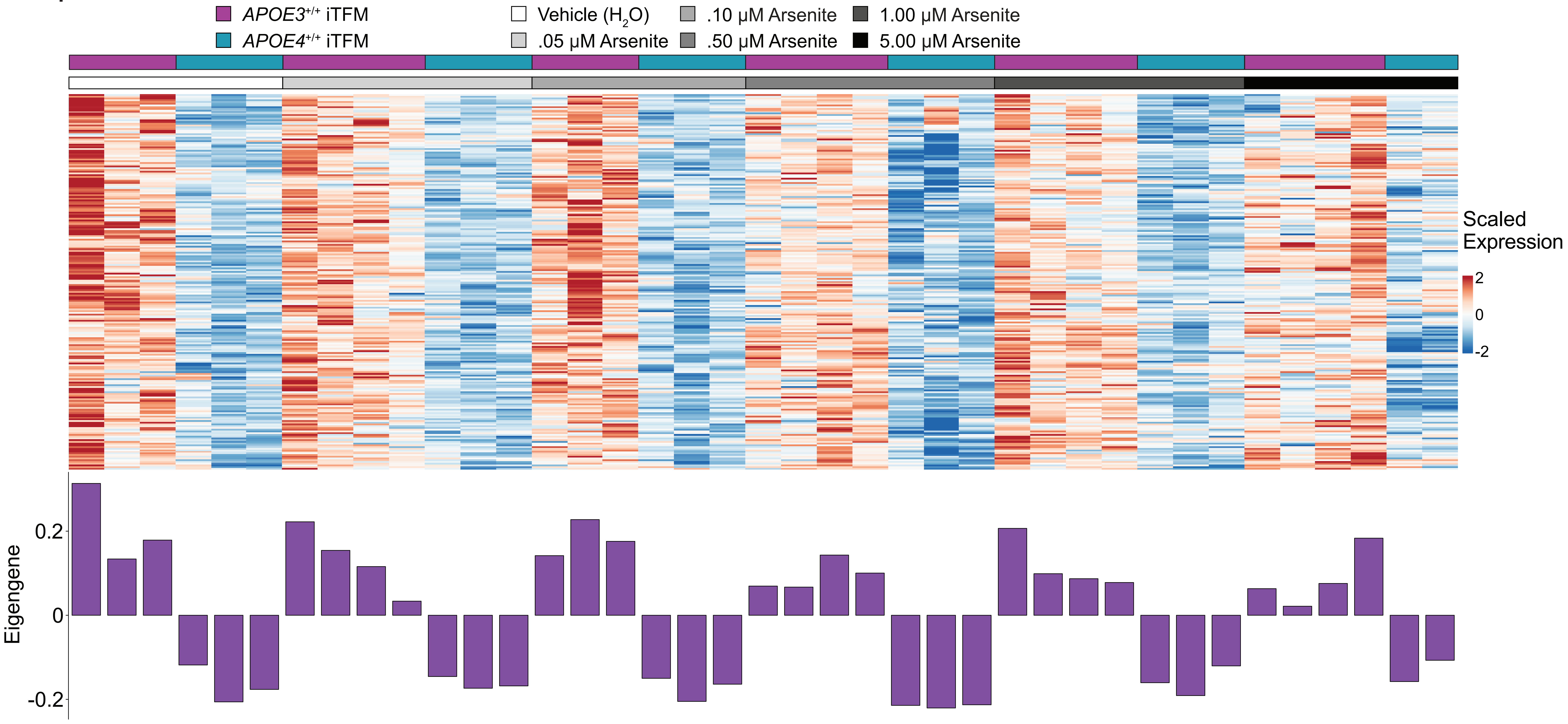

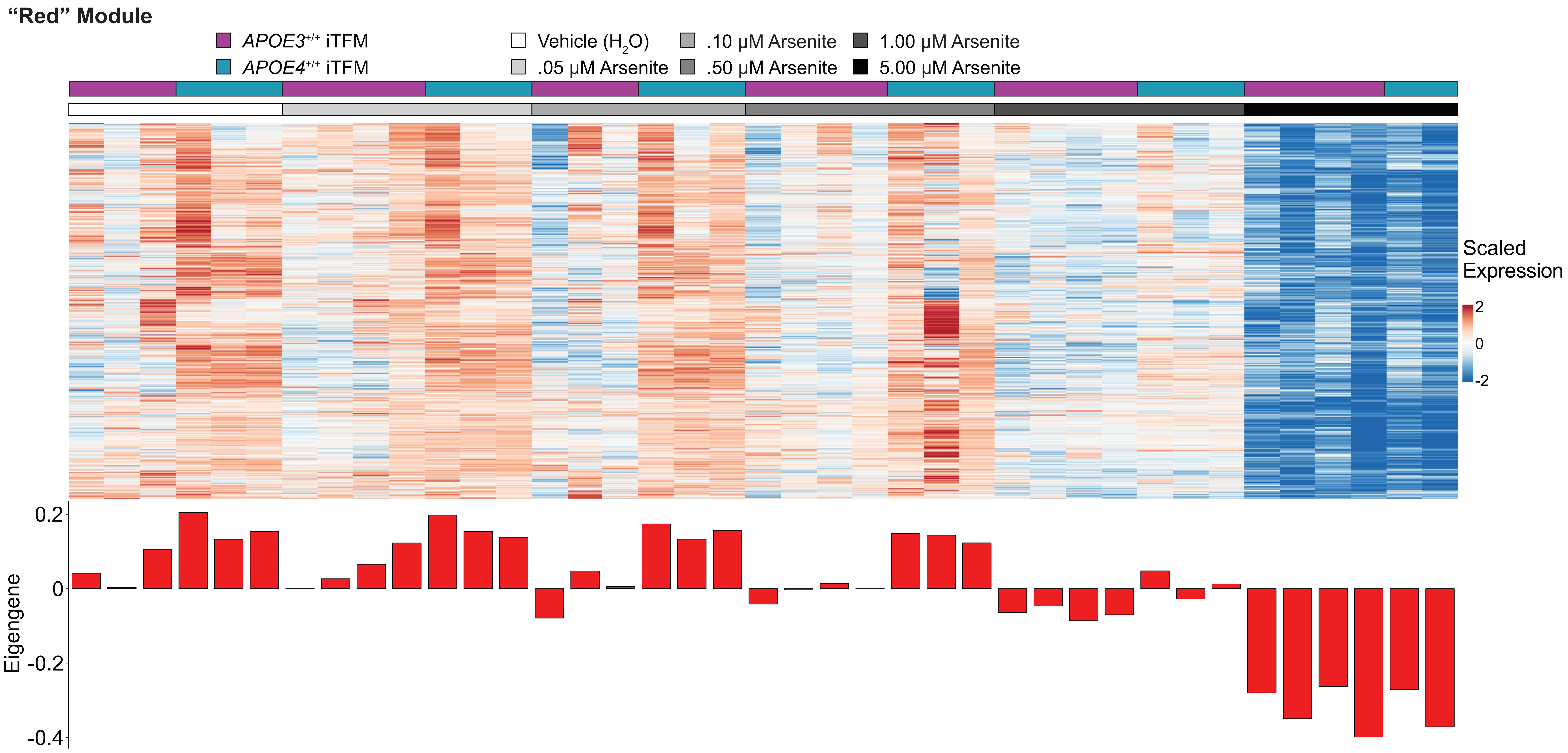
"Red" Module
APOE3+/+ iTFM
APOE4+/+ iTFM
Vehicle (H2O)
.05 μM Arsenite
.10 μM Arsenite
.50 μM Arsenite
1.00 μM Arsenite
5.00 μM Arsenite
Scaled Expression
2
0
-2
Eigengene
0.2
0
-0.2
-0.4

**“Tan” Module**

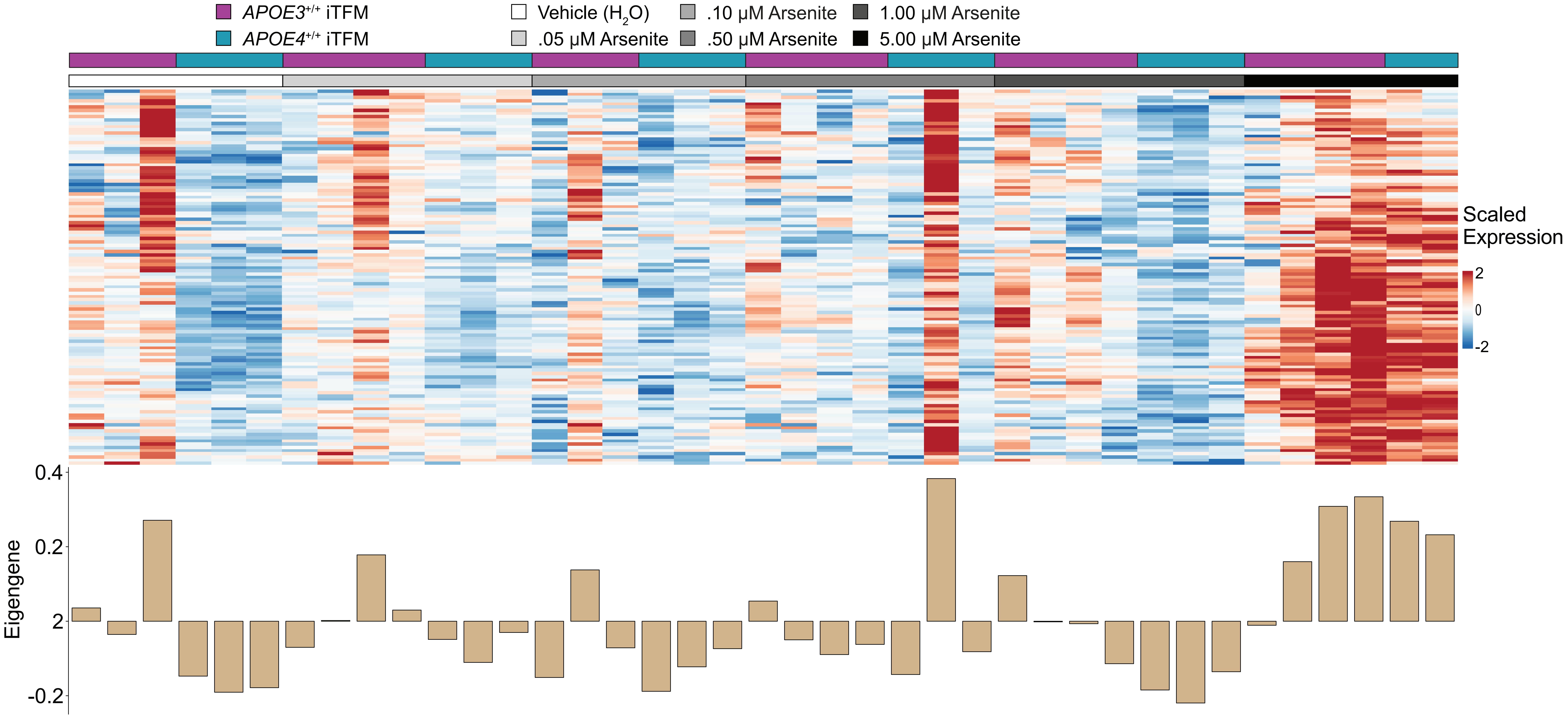

APOE3+/+ iTFM
APOE4+/+ iTFM
Vehicle (H2O)
.05 μM Arsenite
.10 μM Arsenite
.50 μM Arsenite
1.00 μM Arsenite
5.00 μM Arsenite
Scaled Expression
2
0
-2
Eigengene
0.4
0.2
2
-0.2

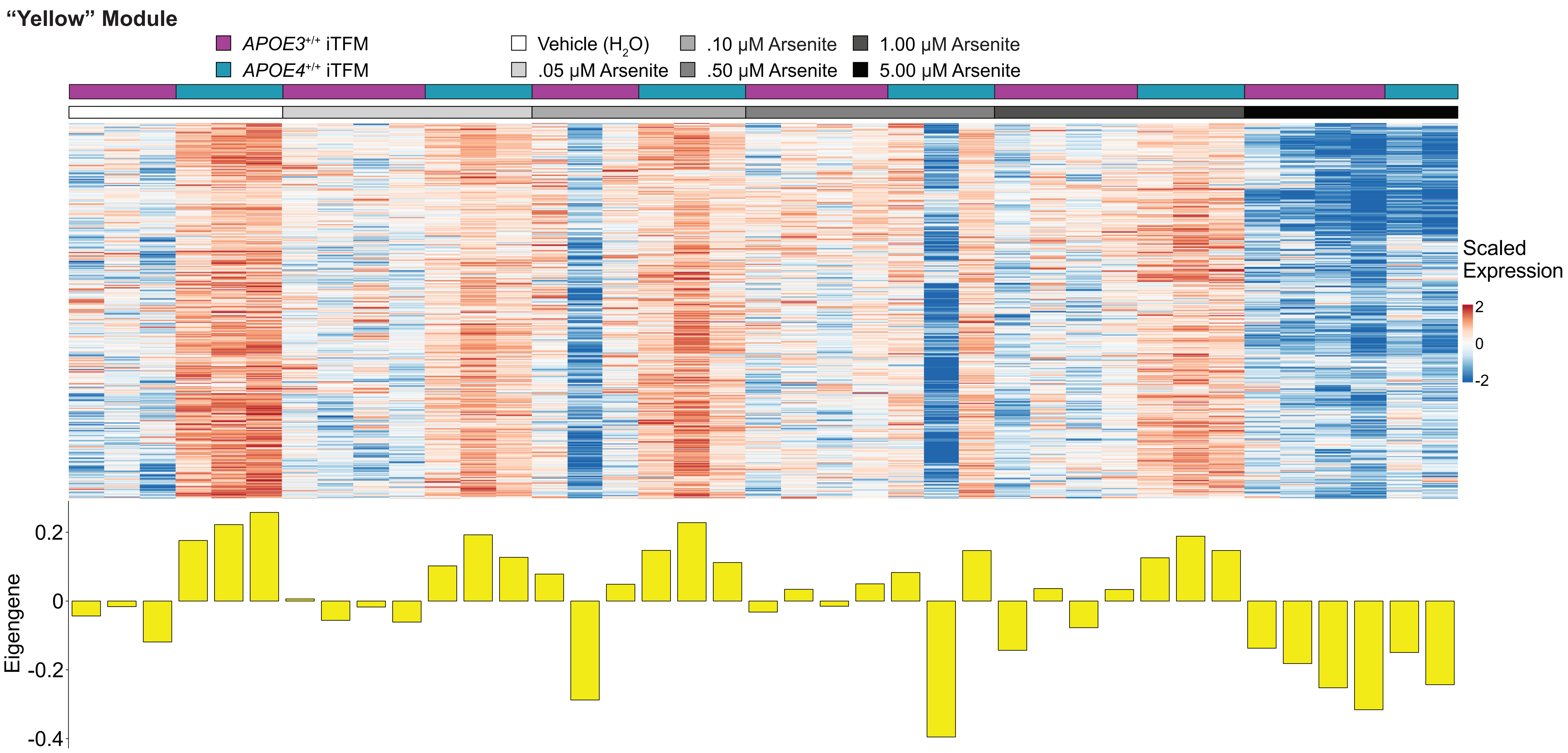
"Yellow" Module
APOE3+/+ iTFM
APOE4+/+ iTFM
Vehicle (H2O)
.05 μM Arsenite
.10 μM Arsenite
.50 μM Arsenite
1.00 μM Arsenite
5.00 μM Arsenite
Scaled Expression
2
0
-2
Eigengene
0.2
0
-0.2
-0.4
